# Regulatory T cells epigenetically reprogrammed from autoreactive effector T cells mitigate established autoimmunity

**DOI:** 10.1101/2025.07.24.665398

**Authors:** Tyler R. Colson, James J. Cameron, Hayley I. Muendlein, Mei-An Nolan, Jamie L. Leiriao, James H. Kim, Alexander N. Poltorak, Xudong Li

**Affiliations:** Graduate Program in Immunology, Tufts Graduate School of Biomedical Sciences, Boston, MA 02111, USA; Department of Immunology, Tufts University School of Medicine; Boston, MA, 02111, USA

## Abstract

Reprogramming autoreactive CD4^+^ effector T (T_eff_) cells into immunosuppressive regulatory T (T_reg_) cells represents a promising strategy for treating established autoimmune diseases. However, the stability and function of such reprogrammed T_regs_ under inflammatory conditions remain unclear. Here, we show that demethylation of core T_reg_ identity genes in T_eff_ cells yields lineage-stable Effector T cell Reprogrammed T_regs_ (ER-T_regs_). A single adoptive transfer of ER-T_regs_ not only prevents autoimmune neuroinflammation in mice when given before disease onset but also arrests its progression when administered after onset. Compared to Foxp3-overexpressing T_eff_ cells, induced T_regs_ from naïve precursors, and endogenous T_regs_, ER-T_regs_ provide superior protection against autoimmune neuroinflammation. This enhanced efficacy stems from their inherited autoantigen specificity and selectively preserved effector-cell transcriptional programs, which together bolster their fitness in inflammatory environments and enhance their suppressive capacity. Our results establish epigenetic reprogramming of autoreactive T_eff_ cells as an effective approach to generate potent, stable T_regs_ for the treatment of refractory autoimmune conditions.

## INTRODUCTION

Regulatory T (T_reg_) cells expressing the transcription factor Foxp3 play an essential role in immune homeostasis by preventing autoimmunity against self-antigens and curtailing deleterious immune responses towards environmental antigens (1, 2). However, naturally occurring endogenous T_regs_ (nT_regs_) are often inadequate at suppressing ongoing inflammation in established autoimmune diseases (3, 4), limiting their therapeutic efficacy (5). This highlights the urgent need to better understand the mechanisms underlying disease-associated T_reg_ functional deficiencies so that novel approaches can be developed to reinvigorate their function for the treatment of autoimmune diseases.

T_reg_ functional deficiency can arise from either inadequate T_reg_ suppressor function on a per-cell basis or a paucity of autoantigen specific T_regs_. The latter may result from impaired differentiation and lineage stability of autoantigen specific T_regs_, leading to their diversion into effector CD4^+^ T (T_eff_) cell fates (6–11). In this regard, reprogramming autoreactive CD4^+^ T_eff_ cells into T_regs_ for adoptive cell therapy presents an attractive option for restoring T_reg_ function. This approach could generate T_regs_ that share the autoantigen specificities of their target T_eff_ cells, potentially enhancing antigen specific suppression (12). However, the development of this approach is hindered by the lack of an effective method to convert CD4^+^ T_eff_ cells into bona fide T_regs_ that exhibit T_reg_-specific gene expression and function (12–14).

The maintenance of T_regs_ cellular identity and suppressive function depends on the stable expression of the transcription factor Foxp3, along with Foxp3-independent core identity genes (15–19). However, whether stable induction of T_reg_ identity genes in committed CD4^+^ T_eff_ cells can establish bona fide T_reg_ identity and suppressive capacity remains unresolved. Pre-existing T_eff_ gene expression may antagonize T_reg_ identity establishment (20), raising concerns that residual effector signatures could destabilize lineage commitment or impair suppressor activity in reprogrammed T_reg_ populations. Alternatively, the inherent T_eff_ gene expression—when combined with the reprogramming process—could actually enhance suppressor function by imparting an effector T_reg_-like gene expression profile, where heightened expression of specific T_eff_ genes correlates with superior suppressive capabilities (21–34). Clarifying these possibilities is crucial not only for advancing this therapeutic strategy but also for providing novel insights into how the core T_reg_ identity gene program interacts with effector gene expression to shape T_reg_ function. Nevertheless, the fundamental question of whether the core T_reg_ identity genes can be stably activated in CD4^+^ T_eff_ cells remains unresolved, hindering deeper exploration of this approach.

T_reg_ gene expression is controlled by intricate epigenetic mechanisms, including DNA methylation and demethylation (11, 35–39). During T_reg_ development, the stable activation of *Foxp3* and other core T_reg_ identity genes requires the demethylation of DNA at cytosine-guanine (CpG) motifs in conserved cis-regulatory regions (16, 40–42), presenting therapeutic targets for boosting T_reg_ function. Indeed, pharmacologically enhancing DNA demethylation in pre-existing T_regs_ augments their fitness and function and accelerates repair of experimental lung injury (43). Importantly, we and others have shown that demethylation at conserved non-coding sequence 2 (CNS2) within the *Foxp3* gene stabilizes its expression in effector T_regs_ by counteracting the transcriptional inhibitory effects of TCR and pro-inflammatory cytokine signaling (44, 45).

These *Foxp3*-inhibitory signals are also highly active in T_eff_ cells (46–48), suggesting that induction of stable *Foxp3* expression in CD4^+^ T_eff_ cells likely requires CNS2 demethylation. Furthermore, we recently demonstrated that CNS2 demethylation requires sustained *Foxp3* transcriptional activation (49), indicating a positive feedback loop where *Foxp3* transcriptional activation and DNA demethylation mutually reinforce each other, facilitating T_reg_ lineage commitment during differentiation. These findings suggest that fostering sustained transcriptional activation of *Foxp3* and other core T_reg_ identity genes in an environment conducive to DNA demethylation may enable epigenetic reprogramming of CD4^+^ T_eff_ cells into T_regs_.

To determine whether autoreactive CD4^+^ T_eff_ cells can be reprogrammed into bona fide T_regs_ for mitigating established autoimmunity, we developed an approach to achieve stable demethylation of T_reg_ identity genes in CD4^+^ T_eff_ cells. Our findings show that this approach generates bona fide T_regs_, which we term Effector T cell Reprogrammed T_regs_ (ER-T_regs_). These ER-T_regs_ exhibit a superior ability to ameliorate established autoimmune neuroinflammation compared to CD4^+^ T_eff_ cells forced to express Foxp3 exogenously, induced T_regs_ (iT_regs_) derived from naïve CD4^+^ T (T_n_) cells, and endogenous nT_regs_. The autoantigen specificity inherited by ER-T_regs_ from autoreactive CD4^+^ T_eff_ cells enables antigen-specific suppression of autoimmune inflammation without compromising normal immune function. Additionally, the selective inheritance of parental T_eff_ gene expression confers ER-T_regs_ superior fitness and suppressor functionality under inflammatory conditions. Thus, demethylation of T_reg_ identity genes in autoreactive CD4^+^ T_eff_ cells can establish a bona fide T_reg_ gene expression program, giving rise to T_regs_ capable of quelling established autoimmune inflammation.

## RESULTS

### Epigenetic reprogramming enables stable induction of *Foxp3* and other core T_reg_ identity genes

*Foxp3* expression is essential for establishing T_reg_ cellular identity (17–19). To induce this identity in pro-inflammatory CD4^+^ T_eff_ cells from mice with experimental autoimmune encephalomyelitis (EAE), we first optimized conditions for efficient *Foxp3* induction. We purified CD4^+^Foxp3-Thy1.1^−^CD44^hi^ T_eff_ cells from *Foxp3^Thy1.1^* reporter mice (50) immunized with myelin oligodendrocyte glycoprotein peptide (MOG) emulsified in complete Freund’s adjuvant (CFA). When activated in vitro under an iT_reg_ differentiation condition consisting of IL-2, TGFβ, and neutralizing antibodies against pro-inflammatory cytokines IL-12, IFNγ, and IL-4 (51, 52), approximately 14% of the T_eff_ cells began expressing Thy1.1 (Figure 1A). Pre-resting T_eff_ cells before activation, along with the addition of retinoic acid (RA) to promote *Foxp3* induction (53–56) and vitamin C (VC) to facilitate DNA demethylation (38, 39), progressively enhanced *Foxp3* induction, resulting in nearly 60% of cells expressing Foxp3 (Figure 1A).

**Figure 1.**
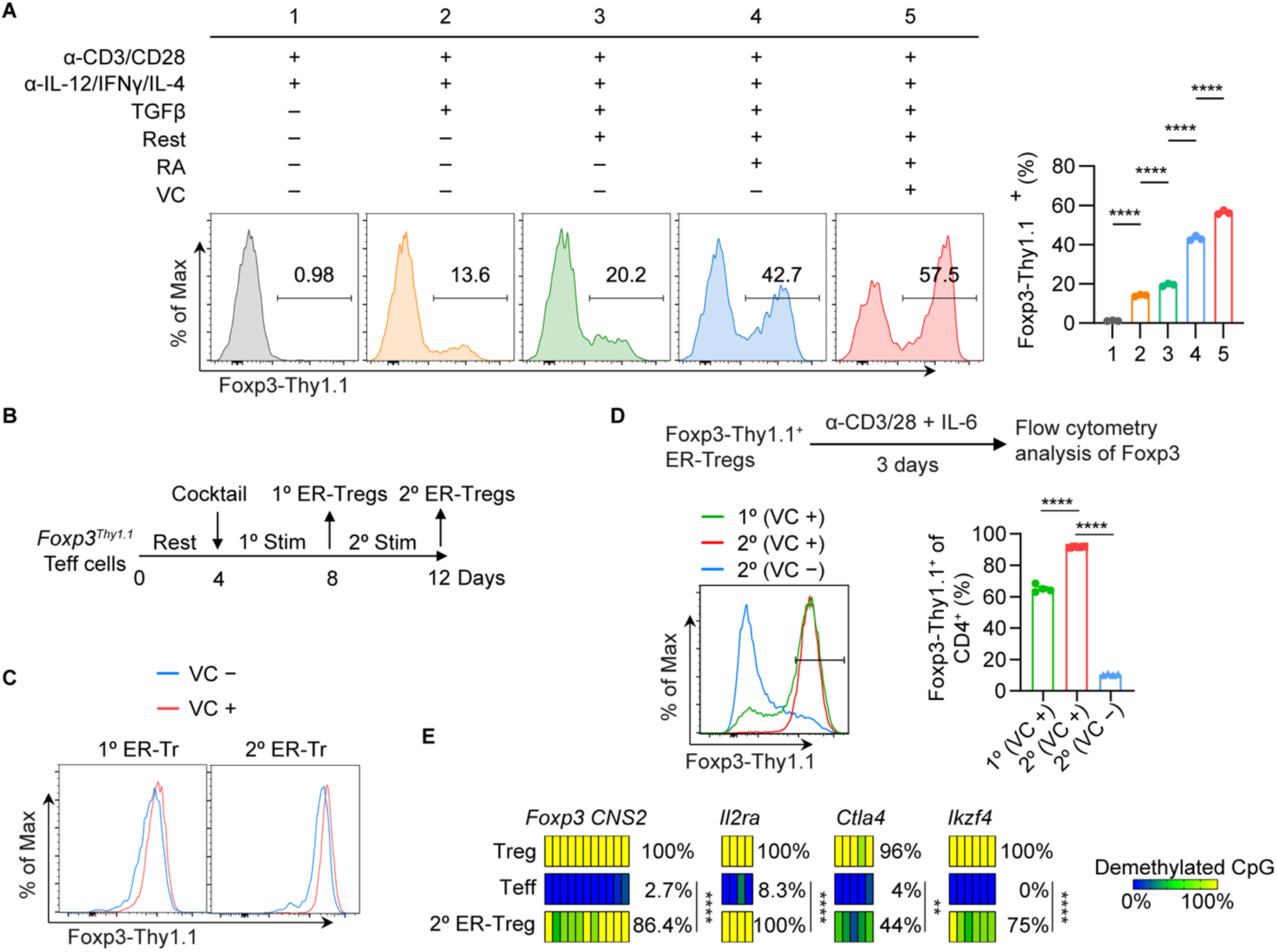
Epigenetic reprogramming of CD4^+^ T_eff_ cells into T_regs_. (**A**) Flow cytometry of Foxp3-Thy1.1 induction in CD4^+^ T_eff_ cells isolated from *Foxp3^Thy1.1^* mice on day 7 following immunization with MOG/CFA and subsequently activated with anti-CD3/CD28 microbeads for 4 days under indicated conditions. Rest indicates resting T_eff_ cells for 4 days prior to their activation. RA indicates retinoic acid. VC indicates vitamin C. (**B**) Schematic of epigenetic reprogramming of CD4^+^ T_eff_ cells into ER-T_regs_. (**C**) Flow cytometry of Foxp3-Thy1.1 expression in 1° and 2° ER-T_regs_ generated in the presence or absence of VC. (**D**) Flow cytometry of Foxp3-Thy1.1 expression in 1° and 2° ER-T_regs_ generated in the presence or absence of VC and subsequently re-stimulated for 3 days in the presence of IL-6. (**E**) Heatmaps of CpG demethylation patterns at specific loci in indicated cell types, analyzed with bisulfite-sequencing. Each bar represents a CpG site. Mean ± SEM. **p < 0.01, ****p < 0.0001, one-way ANOVA and Holm-Šídák test in (**A** and **D**), and two-way ANOVA in (**E**).

Sustained transcriptional activation of *Foxp3* enhances its epigenetic stabilization by promoting CNS2 demethylation (49). To test whether maintaining *Foxp3* activation in ER-T_regs_ through re-stimulation with the ER-T_reg_ reprogramming cocktail improves *Foxp3* stability, we re-stimulated the cells in this context (Figure 1B). Both re-stimulation and the inclusion of VC during reprogramming increased *Foxp3* expression (Figure 1C), and more importantly, improved its stability when ER-T_regs_ were subsequently exposed to the pro-inflammatory cytokine IL-6 (Figure 1D). Notably, restimulated ER-T_regs_ displayed significant DNA demethylation at T_reg_-specific demethylation regions, including *Foxp3 CNS2*, *Il2ra*, *Ctla4*, and *Ikzf4* (16), compared to parental T_eff_ cells (Figure 1E). Collectively, these findings suggest that both features defining T_reg_ cellular identity – stable *Foxp3* expression and epigenetic activation of core T_reg_ identity genes – can be successfully established in committed CD4^+^ T_eff_ cells by epigenetic reprogramming.

### Adoptive transfer of ER-T_regs_ prevents EAE development and ameliorates established EAE

To assess the therapeutic potential of ER-T_regs_ in curbing autoimmune inflammation, we adoptively transferred ER-T_regs_ derived from MOG/CFA-primed CD4^+^ T_eff_ cells into CD45 congenically distinct mice one day before inducing EAE via MOG/CFA immunization and pertussis toxin (PT) injections. Remarkably, ER-T_reg_ transfer nearly abolished clinical manifestations of EAE, in stark contrast to the rapid disease progression observed in control animals (Figure 2A). Consistent with this protection, CD4^+^ T cell infiltration into the spinal cord was significantly reduced in ER-T_regs_-treated mice (Figure 2B). Approximately 90% of transferred ER-T_regs_ maintained Foxp3 expression, demonstrating robust in vivo stability under inflammatory conditions (Figure 2C). Although ER-T_regs_ comprised only about 2% of all T_regs_ in the draining lymph nodes (Supplemental Figure 1A), they exhibited a significantly higher frequency of RORγt^+^ populations and a trending increase in RORγt^+^c-MAF^+^ populations compared to endogenous T_regs_ (Supplemental Figure 1B). This distinct phenotypic profile, combined with their scarcity, collectively suggests that ER-T_regs_ possess enhanced per-cell suppressive potency compared to endogenous T_regs_ under inflammatory conditions.

**Figure 2.**
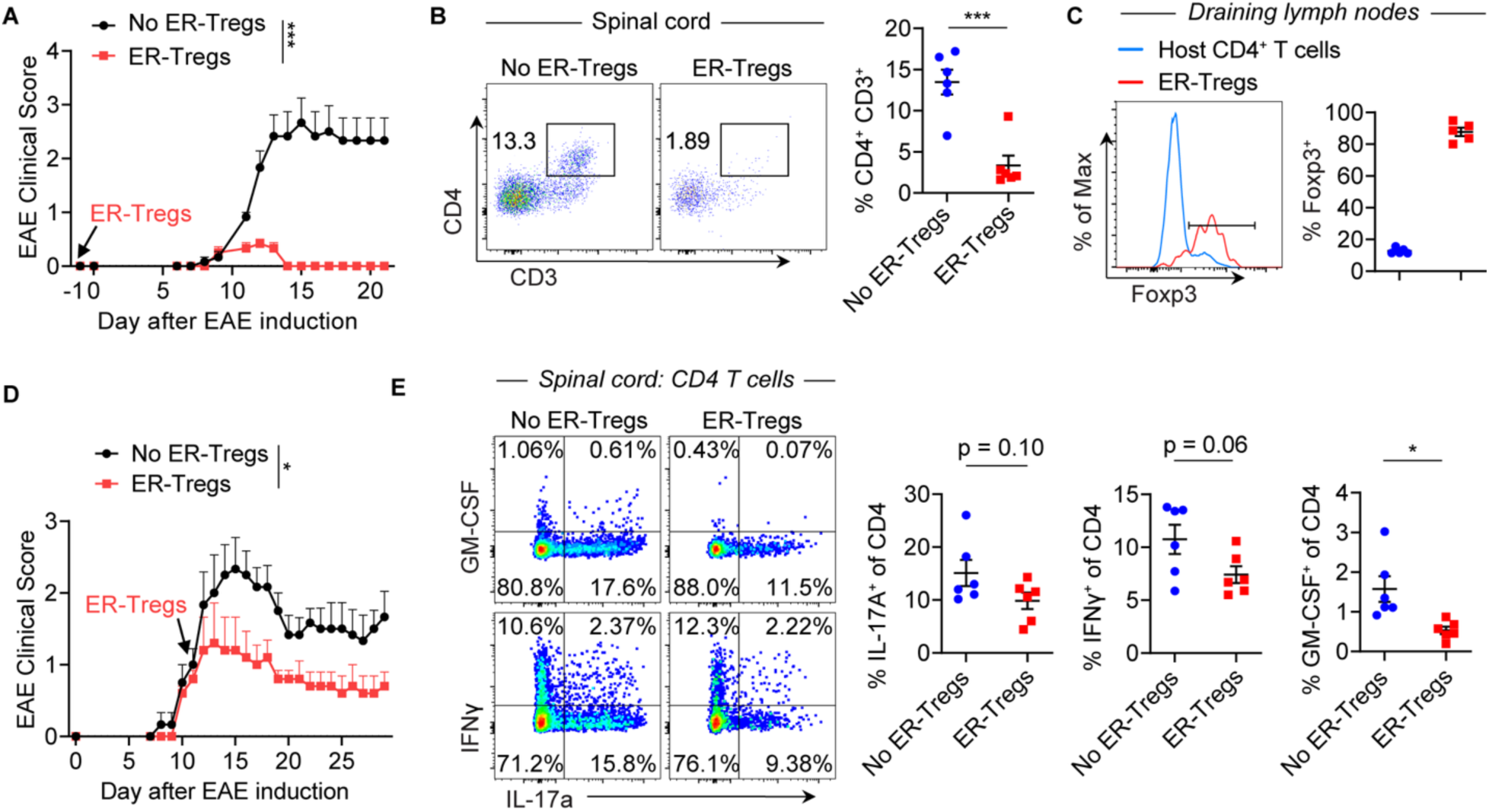
Adoptive transfer of ER-T_regs_ prevents EAE development and ameliorates established EAE. (**A-C**) EAE was induced via MOG/CFA immunization in CD45.2^+^ mice with or without adoptive transfer of ER-T_regs_ reprogrammed from MOG/CFA-primed CD45.1^+^*Foxp3^Thy1.1^*CD4^+^ T_eff_ cells, administered one day prior to immunization. Flow cytometry analyses were conducted at 21 days post-immunization (dpi). n = 6 per group. Data are representative of two independent experiments. (**A**) EAE disease curve. (**B**) Flow cytometry analysis of the frequencies of spinal cord CD4^+^ T cells. (**C**) Flow cytometry of Foxp3 expression in host CD4^+^ T cells and transferred ER-T_regs_ within the draining lymph nodes (LNs) of mice that received ER-T_regs_. (**D and E**) EAE was induced via MOG/CFA immunization in CD45.2^+^ mice with or without adoptive transfer of ER-T_regs_ reprogrammed from MOG/CFA-primed CD45.1^+^*Foxp3^Thy1.1^*T_eff_ cells, administered at 11 dpi. Flow cytometry analyses were conducted at 29 dpi. n = 6 per group. Data are representative of two independent experiments. (**D**) EAE disease curve. (**E**) Flow cytometry of IFNγ and GM-CSF expression in CD4^+^ T cells in spinal cord. Mean ± SEM. *p < 0.05, ***p < 0.001, unpaired two-sided t-test of Area under the curve (AUC) in (**A** and **D**) and unpaired two-sided t-test in (**B** and **E**).

To assess the suppressive capacity of ER-T_regs_ in established EAE, we adoptively transferred ER-T_regs_ derived from MOG/CFA-primed CD4^+^ T_eff_ cells into CD45 congenically distinct mice at disease onset (clinical score ∼1). ER-T_reg_ transfer attenuated disease progression (Figure 2D) and significantly reduced spinal cord infiltration by GM-CSF-producing CD4^+^ T_eff_ cells (Figure 2E), a key driver of neuroinflammation (57–59). Detectable ER-T_regs_ were observed in only two recipients, likely reflecting the contraction of the transferred population as inflammation resolved. In these mice, ER-T_regs_ accounted for approximately 20% of spinal cord Foxp3^+^ cells (Supplemental Figure 1C). Strikingly, spinal cord ER-T_regs_ exhibited a higher frequency of RORγt^+^ and RORγt^+^c-MAF^+^ subsets compared to endogenous T_regs_ (Supplemental Figure 1D), further underscoring their distinct phenotype. Notably, ER-T_regs_ lacked expression of inflammatory cytokines IFNγ or GM-CSF, although about 20% produced IL-17A (Supplemental Figure 1E). Collectively, these findings demonstrate that ER-T_regs_ retain robust lineage stability and suppressive potency in vivo, even within an established inflammatory niche, with a single transfer sufficient to ameliorate ongoing autoimmune pathology.

### Foxp3 expression is required but not sufficient for ER-T_reg_ suppressor function

While Foxp3 expression is essential for the development and function of most endogenous T_regs_ (17–19), a recent study demonstrated that Foxp3 is dispensable for the fitness of microbiota-dependent peripherally induced T_regs_ (pT_regs_) and their ability to suppress colonic T cell expansion (60). To determine whether *Foxp3* activation is necessary for ER-T_reg_ suppressor function, we used CRISPR/Cas9 to ablate *Foxp3* in *Foxp3^Thy1.1^R26^Cas9^* ER-T_regs_. We transduced these cells with a retroviral vector expressing a single guide RNA targeting *Foxp3* (sgFoxp3) and compared their capacity to suppress GM-CSF expression in MOG/CFA-primed responder CD4^+^ T_eff_ cells to that of ER-T_regs_ transduced with a non-targeting sgRNA (sgNT). *Foxp3* ablation completely abolished ER-T_reg_-mediated suppression of GM-CSF in CD4^+^ T_eff_ cells following MOG stimulation (Figure 3A). Moreover, Foxp3-deficient ER-T_regs_ exhibited increased IL-17A expression (Figure 3B), consistent with studies showing that Foxp3 is critical for repressing IL-17A in pT_regs_ (60). In addition, *Foxp3* ablation led to reduced IL-10 expression in ER-T_regs_ (Figure 3B). Collectively, these results indicate that activation of *Foxp3* is indispensable for the suppressive function of ER-T_regs_.

**Figure 3.**
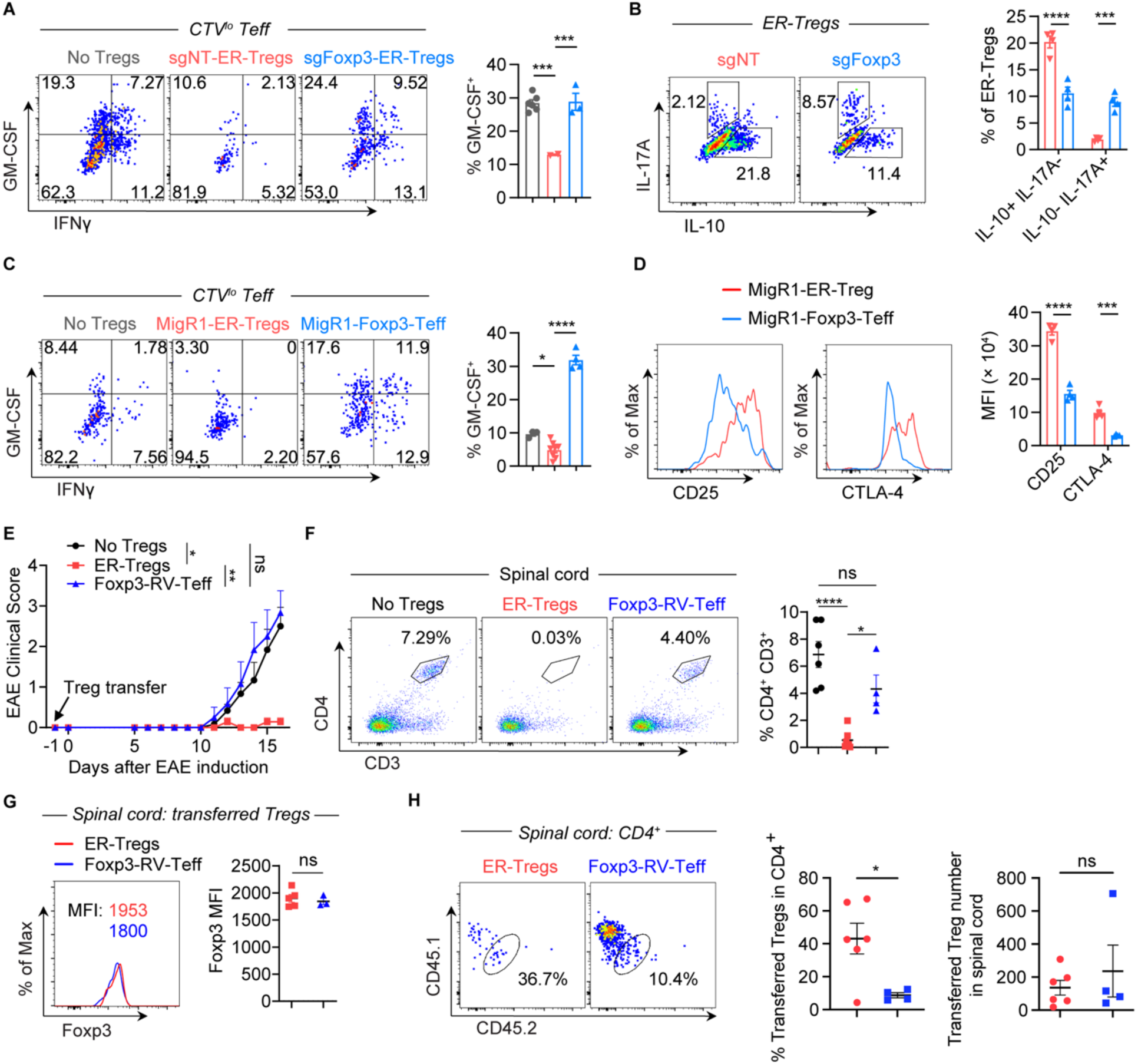
Foxp3 expression is required but not sufficient for the suppressor function of ER-T_regs_. (**A**) Flow cytometry of IFNγ and GM-CSF expression in CTV^lo^ CD4^+^ T_eff_ cells, cocultured for 3 days with T-cell-depleted splenocytes serving as antigen presenting cells (APCs), in the presence of MOG and the presence or absence of *Foxp3^Thy1.1^ R26^Cas9^* ER-T_regs_ transduced with the retroviral vector (RV) expressing single guide RNA targeting Foxp3 (sgFoxp3) or the non-targeting sgRNA (sgNT). (**B**) Flow cytometry of IL-10 and IL-17A expression in ER-T_regs_ as in (**A**). (**C**) Flow cytometry of IFNγ and GM-CSF expression in CTV^lo^ CD4^+^ T_eff_ cells, cocultured for 3 days with APCs, in the presence of MOG and the presence or absence of Foxp3^Thy1.1^ ER-T_regs_ transduced with MigR1 empty vector or CD4^+^ T_eff_ cells forced to express Foxp3 via retroviral transduction with MigR1-Foxp3. (**D**) Flow cytometry of CD25 and CTLA-4 expression in MigR1-ER-T_regs_ and MigR1-Foxp3 CD4^+^ T_eff_ cells as in (**C**). (**E-G**) EAE was induced via MOG/CFA immunization in CD45.2^+^ mice with or without adoptive transfer of CD45.1^+^*Foxp3^Thy1.1^*MOG/CFA-primed CD4^+^ T_eff_ cells reprogrammed into ER-T_regs_ or forced to express Foxp3 via retroviral transduction (Foxp3-RV-T_eff_), administered one day prior to immunization. Flow cytometry analyses were conducted at 16 dpi. n = 6 per group. (**E**) EAE disease curve. (**F**) Flow cytometry analysis of the frequencies of spinal cord CD4^+^ T cells. (**G**) Flow cytometry of Foxp3 expression in adoptively transferred ER-T_regs_ and Foxp3-RV CD4^+^ T_eff_ cells in the spinal cord. (**H**) Flow cytometry of the frequencies and numbers of ER-T_regs_ or FRV-T_regs_ in the spinal cord. Mean ± SEM. *p < 0.05, **p < 0.01, ***p < 0.001, ****p < 0.0001, one-way ANOVA and Holm-Šídák test in (**A**, **C**, **E**, and **F**) and unpaired two-sided t-test in (**B**, **D**, **G,** and **H**).

To investigate whether epigenetic activation of Foxp3-independent T_reg_ identity genes (Figure 1E) is essential for ER-T_reg_ suppressive function, we forced Foxp3 expression in CD4^+^ T_eff_ cells via retroviral transduction and assessed their regulatory capacity. In contrast to ER-T_regs_, Foxp3-expressing T_eff_ cells failed to suppress GM-CSF production in responder CD4^+^ T_eff_ cells upon MOG stimulation. Instead, they amplified inflammatory responses, likely due to their inherently elevated expression of pro-inflammatory cytokines (Figure 3C; Supplemental Figure 2A). Furthermore, these cells exhibited markedly reduced expression of critical T_reg_ effector molecules, including CD25 and CTLA-4, which are encoded by TSDR-containing *Il2ra* and *Ctla4*, respectively (61, 62) (Figure 3D).

To evaluate the in vivo relevance of these findings, we compared the therapeutic efficacy of Foxp3-expressing T_eff_ cells with that of ER-T_regs_ in EAE. Adoptive transfer of Foxp3-expressing T_eff_ cells neither attenuated disease progression nor reduced CD4^+^ T cell infiltration in the spinal cord (Figure 3E-F). Although these cells expressed Foxp3 at levels comparable to ER-T_regs_ (Figure 3G), their relative abundance in the spinal cord was significantly lower (Figure 3H). These results underscore the necessity of Foxp3-independent epigenetic reprogramming for effective ER-T_reg_ functionality.

To evaluate differences between Foxp3-expressing T_eff_ cells and ER-T_regs_, we directly compared their fitness and phenotype in immunocompetent EAE hosts. Foxp3-expressing T_eff_ cells displayed reduced splenic engraftment, lower proliferation (as indicated by Ki-67 staining), and diminished expression of Helios and CD25 compared to ER-T_regs_ (Supplemental Figure 2B-F). Given that Helios (encoded by *Ikzf2*) promotes T_reg_ stability and survival (63, 64) and CD25 (encoded by *Il2ra*) enhances IL-2–dependent survival and function (62, 65, 66), the diminished suppressive capacity of Foxp3-expressing T_eff_ cells thus likely results from inadequate epigenetic priming of critical genes (e.g., *Helios*, *Il2ra*), thereby compromising their resilience in inflammatory environments.

### Inherited autoantigen specificity confers superior functionality to ER-T_regs_

Antigen specificity is critical for T_reg_ suppressor function, suggesting that the inheritance of parental T_eff_ autoantigen specificity may enhance ER-T_reg_ activity. To investigate this possibility, we examined whether the pro-inflammatory environment during autoimmune inflammation hinders the de novo differentiation of MOG-specific T_regs_ in EAE. We adoptively transferred CellTrace Violet (CTV)-labeled Foxp3-Thy1.1^−^ conventional CD4^+^ T (T_conv_) cells or Foxp3-Thy1.1^+^ nT_regs_ from unimmunized donor mice into CD45 congenically distinct mice, followed by immunization with MOG/CFA. Fewer than 1% of CTV^lo^ donor T_conv_ cells expressed Foxp3-Thy1.1, whereas the majority of CTV^lo^ donor nT_regs_ maintained Foxp3 expression (Figure 4A), indicating that de novo differentiation of T_conv_ cells into T_regs_ does not occur in EAE. Together, these findings suggest that the pro-inflammatory environment in EAE drives MOG-specific CD4^+^ naïve T cells to differentiate into T_eff_ cells rather than T_regs_.

**Figure 4.**
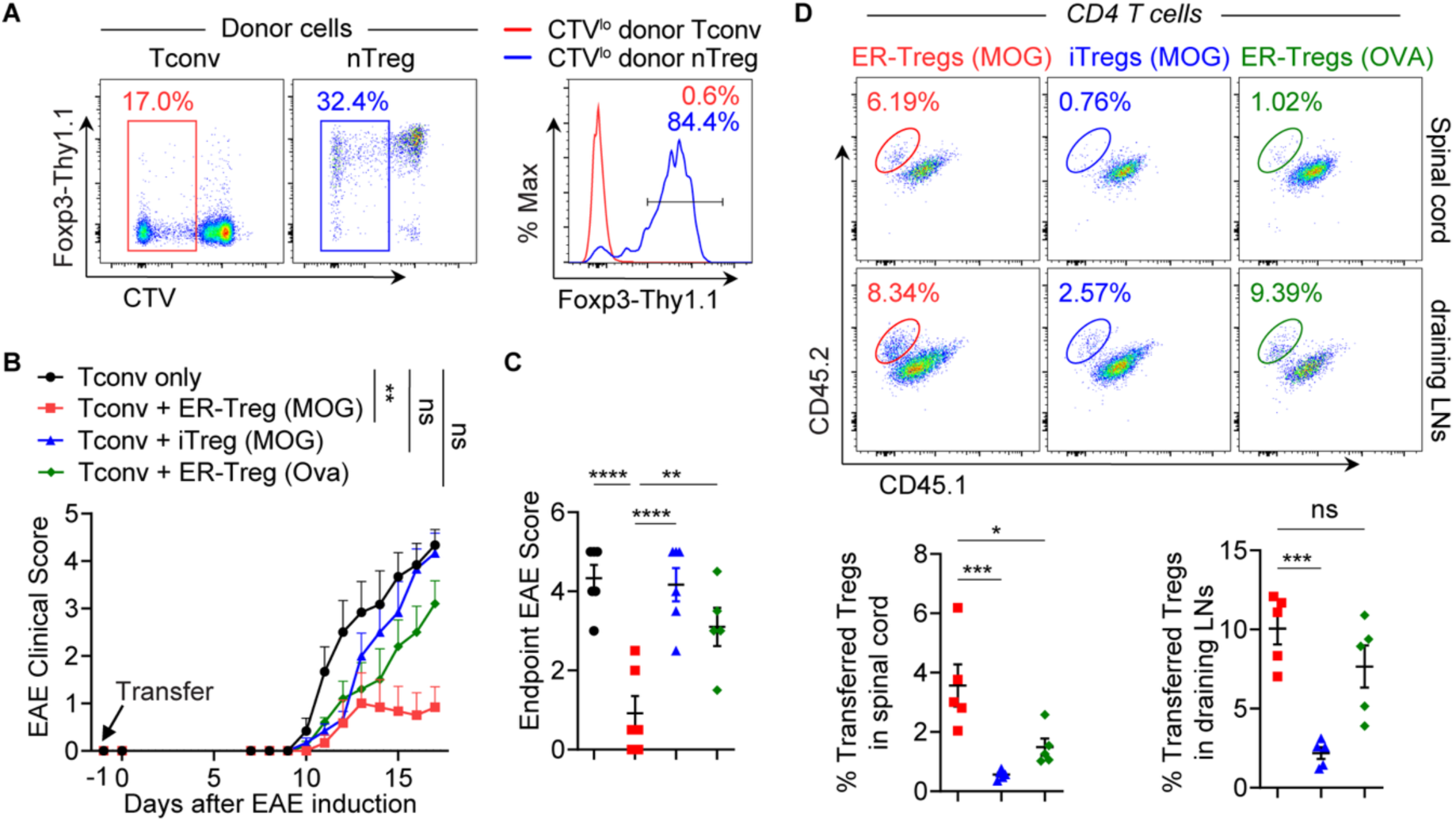
Inheritance of autoantigen specificity contributes to superior suppressive function of ER-T_regs_ as compared to induced T_regs_. (**A**) Flow cytometry of Foxp3 expression in CTV^lo^ CD4^+^ T_conv_ and nT_reg_ cells 8 days after they were isolated from CD45.2^+^*Foxp3^Thy1.1^* mice, labeled with CTV, and adoptively transferred into CD45.1^+^*Foxp3^Thy1.1^*mice, which were subsequently immunized with MOG/CFA one day after adoptive transfer. (**B-D**) EAE was induced in *Rag1^−/−^* mice via MOG/CFA immunization one day after adoptive transfer of MOG/CFA-primed CD4^+^ T_conv_ cells with or without co-transfer of congenically distinct ER-T_regs_ reprogrammed from MOG/CFA- or OVA/CFA-primed CD4^+^ T_eff_ cells, or co-transfer of induced T_regs_ (iT_regs_) generated with in vitro differentiation of T_n_ cells isolated from MOG/CFA-primed mice. Flow cytometry analyses were conducted at 17 dpi. n = 6 per group. (**B**) EAE disease curve. (**C**) EAE scores at 17 dpi. (**D**) Flow cytometry analysis of the frequencies of adoptively transferred ER-T_regs_ or iT_regs_ (CD45.1^−^CD45.2^+^) in the spinal cord and draining LNs. Mean ± SEM. *p < 0.05, **p < 0.01, ***p < 0.001, ****p < 0.0001, unpaired two-sided t-test in (**B**), one-way ANOVA and Holm-Šídák test in (**C, D**).

To assess whether inherited myelin autoantigen specificity contributes to ER-T_reg_ suppressor function, we performed adoptive transfers into *Rag1^-/-^* mice. Specifically, we transferred MOG/CFA-primed CD4^+^ T_conv_ cells alone or co-transferred them with CD45 congenically distinct ER-T_regs_ reprogrammed from CD4^+^ T_eff_ cells primed in vivo with either MOG/CFA or Ovalbumin peptide (OVA) emulsified in CFA. We also included iT_regs_ differentiated from T_n_ cells from MOG/CFA-immunized mice. Following recipient immunization with MOG/CFA and PT administration, mice receiving only T_conv_ cells developed severe EAE (Figure 4, B and C). In contrast, co-transfer of ER-T_regs_ derived from MOG/CFA-primed CD4^+^ T_eff_ cells substantially mitigated EAE, whereas co-transfer of ER-T_regs_ derived from OVA/CFA-primed CD4^+^ T_eff_ cells or iT_regs_ did not. Notably, the suppressive efficacy of the transferred T_regs_ correlated positively with their frequencies in the spinal cord (Figure 4D). These findings suggest that inherited myelin autoantigen specificity contributes to the ability of ER-T_regs_ to curtail EAE development, at least in part by enhancing their ability to accrue in the inflamed tissue. Conversely, the diminished suppressive capacity of autologous iT_regs_ may reflect diminished autoantigen specificity, as inflammatory conditions preferentially drive the differentiation of autoreactive T_n_ cells into T_eff_ cells.

### ER-T_regs_ suppress EAE in an autoantigen-specific manner without inhibiting immune response against a non-myelin foreign antigen

Initial adoptive transfer experiments revealed that both iT_regs_ and OVA-specific ER-T_regs_ transiently delayed EAE progression at early stages (Figure 4B), suggesting the possibility of antigen-nonspecific immunosuppressive effects. To clarify this, we developed a refined reprogramming protocol leveraging MOG-induced CTV dilution to isolate MOG-specific from MOG-nonspecific ER-T_reg_ populations. Transfer of MOG-specific ER-T_regs_, but not their nonspecific counterparts, significantly attenuated EAE severity and reduced spinal cord infiltration by GM-CSF^+^ CD4^+^ T_eff_ cells (Figure 5, A and B). Moreover, MOG-specific ER-T_regs_ exhibited enhanced tissue fitness, as evidenced by substantially higher frequencies and total numbers in the spinal cord (Figure 5C), despite comparable splenic engraftment (Supplemental Figure 3A). They also demonstrated superior lineage stability, with elevated frequencies of Foxp3-Thy1.1^+^ cells and higher Thy1.1 MFI, as well as enrichment for c-MAF^+^RORγt^+^ subsets (Supplemental Figure 3B–C), a phenotype linked to enhanced Th17 suppression (31–34). These findings align with previous studies showing that TCR activation enhances Foxp3 expression, functional specialization, and tissue homing in T_regs_ (67, 68), and underscores the necessity of myelin antigen specificity for ER-T_regs_ to durably suppress CNS inflammation.

**Figure 5.**
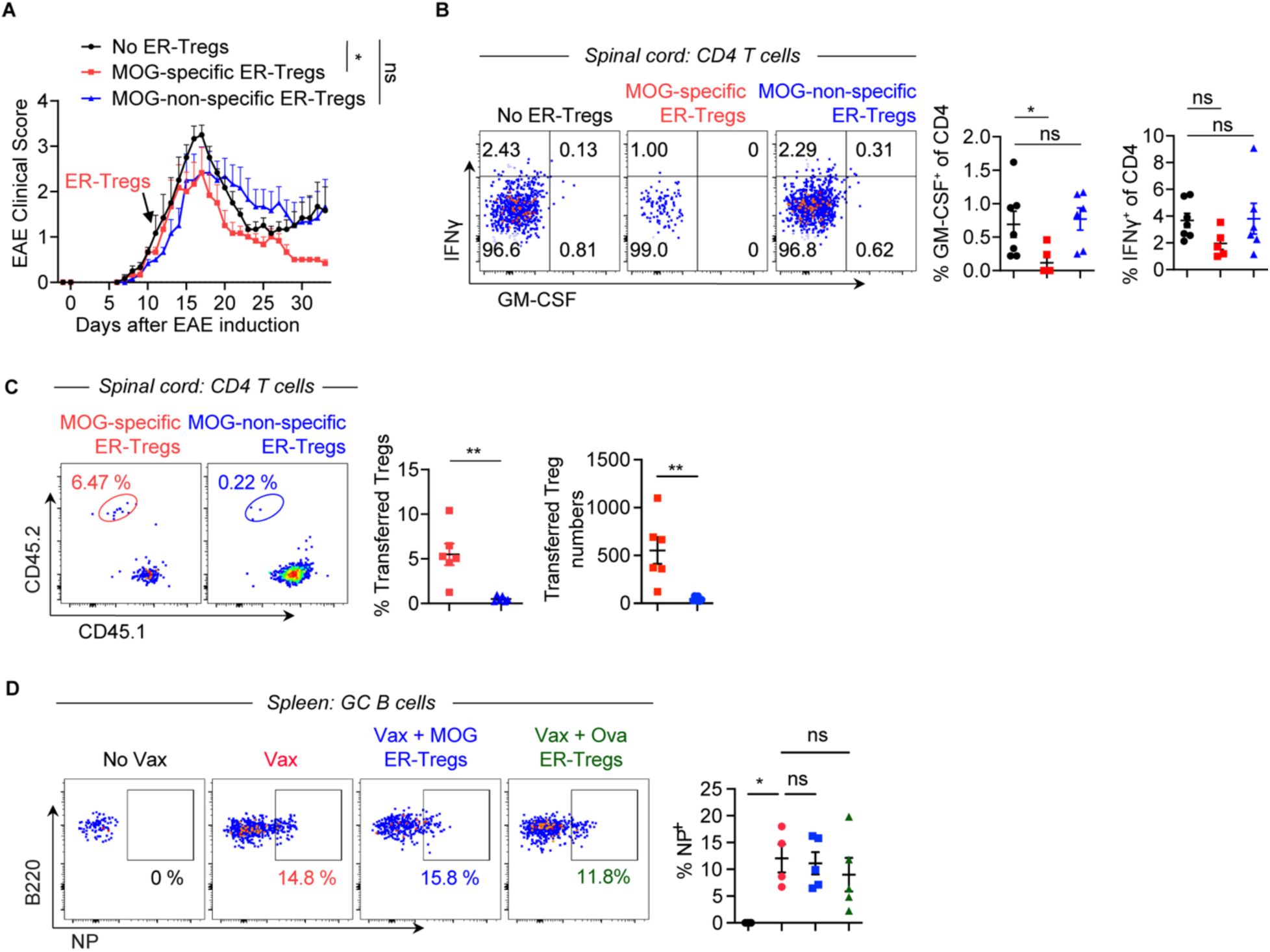
ER-T_regs_ confer autoantigen-specific suppression of EAE without compromising vaccine-elicited immune responses against a foreign antigen. (**A-C**) EAE was induced via MOG/CFA immunization in CD45.1^+^ mice with or without adoptive transfer of CD45.2^+^*Foxp3^Thy1.1^* MOG-specific or MOG-non-specific ER-T_regs_, administered at 11 dpi. Flow cytometry analyses were conducted at 33 dpi. n = 6 per group. (**A**) EAE disease curve. (**B**) Flow cytometry of IFNγ and GM-CSF expression in CD4^+^ T cells in the spinal cord. (**C**) Flow cytometry analysis of the frequencies and total numbers of adoptively transferred ER-T_regs_ (CD45.1^−^CD45.2^+^) in the spinal cord. (**D**) Flow cytometry analysis of the frequencies of NP-specific (NP-PE^+^) germinal center B cells in the spleens of mice 11 days post-immunization with NP-OVA/Alum with or without adoptive transfer of MOG-specific or OVA-specific ER-T_regs_ administered one day prior to immunization. Mean ± SEM. *p < 0.05, **p < 0.01, one-way ANOVA and Holm-Šídák test in (**A**, **B**, and **D**), and unpaired two-sided t-test in (**C**).

To evaluate antigen-nonspecific immunosuppression, we transferred ER-T_regs_ generated from either CFA/MOG- or CFA/Ova-primed T_eff_ cells into mice immunized with nitrophenol-conjugated ovalbumin (NP-Ova) in Alum. Neither MOG-specific nor OVA-specific ER-T_regs_ significantly inhibited NP-specific germinal center B cell responses (Figure 5D), although OVA-specific ER-T_regs_ exhibited a modest, non-significant trend toward suppression. These results suggest that ER-T_reg_-mediated suppression is tightly restricted to their cognate antigen and relies on the inflammatory context.

### ER-T_regs_ selectively inherit parental T_eff_ gene expression

To determine whether parental T_eff_ gene expression confers a distinct transcriptional profile to ER-T_regs_, we performed bulk RNA-seq on ER-T_regs_, nT_regs_, and CD4^+^ T_eff_ cells—all isolated from MOG/CFA-immunized mice to ensure uniform in vivo exposure. Principal component analysis revealed that the ER-T_reg_ transcriptome more closely resembles that of nT_regs_ than CD4^+^ T_eff_ cells (Figure 6A). Moreover, ER-T_regs_ expressed significantly higher levels of core T_reg_ identity genes, including *Foxp3*, *Itgae*, *Il2ra*, *Ctla4*, and *Ikzf4* (22), compared to CD4^+^ T_eff_ cells (Figure 6B). Gene set enrichment analysis (GSEA) further demonstrated that genes typically upregulated (or downregulated) in T_regs_ relative to CD4^+^ T_conv_ cells are similarly upregulated (or downregulated) in ER-T_regs_ relative to CD4^+^ T_eff_ cells (Figure 6C), confirming the successful establishment of a T_reg_ gene expression program in ER-T_regs_.

**Figure 6.**
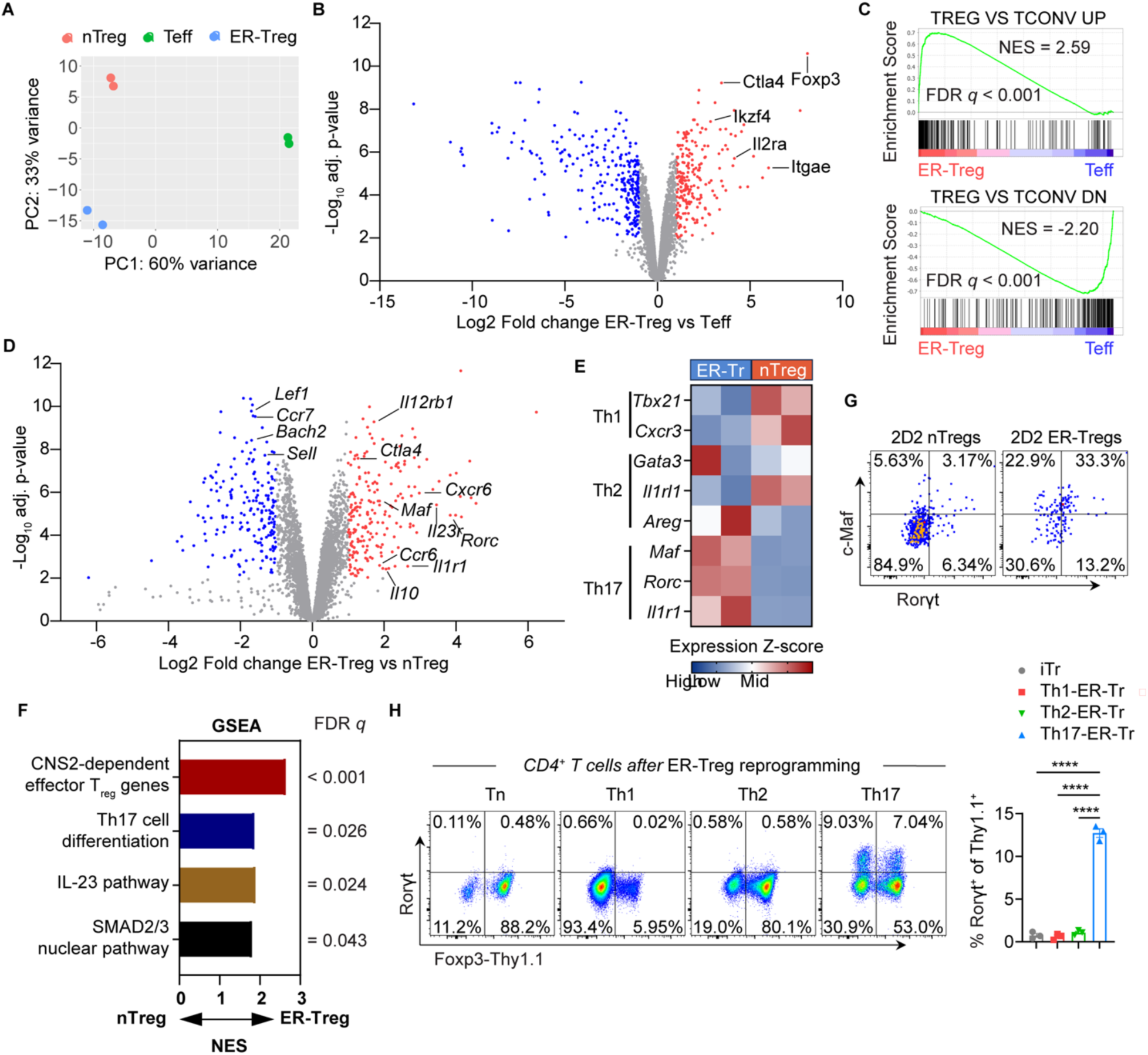
ER-T_regs_ exhibit elevated expression of select parental T_eff_ genes. (**A-F**) RNA-seq analysis of the transcriptomes of ER-T_regs_, nT_regs_, and CD4^+^ T_eff_ cells. CD45.1^+^*Foxp3^Thy1.1^*ER-T_regs_ were adoptively transferred into CD45.2^+^*Foxp3^Thy1.1^* mice one day prior to MOG/CFA immunization. The transcriptomes of transferred ER-T_regs_ and host nT_reg_ and CD4^+^ T_eff_ cells were determined by bulk RNA-seq at 7 dpi. (**A**) Principal component analysis of ER-T_reg_, nT_reg_, and CD4^+^ T_eff_ transcriptomes. (**B**) Volcano plot showing the differential expression of genes between ER-T_regs_ and T_eff_ cells. (**C**) Gene Set Enrichment Analysis (GSEA) of the expression of T_reg_-specific genes in ER-T_regs_ and nT_regs_. (**D**) Volcano plot showing the differential expression of genes between ER-T_regs_ and nT_regs_. (**E**) Normalized gene expression levels for selected lists of T helper genes in ER-T_regs_ and nT_regs_. (**F**) GSEA of the expression of indicated gene sets in ER-T_regs_ and nT_regs_. (**G**) Flow cytometry of c-Maf and Rorγt expression in MOG/CFA-primed 2D2 nT_regs_ and ER-T_regs_ following in vitro activation in the presence of IL-2 for 3 days. (**H**) Flow cytometry of Rorγt and Foxp3-Thy1.1 expression in CD4^+^ T_n_ cells and in vitro differentiated T helper cells following their 1° stimulation under the ER-T_reg_ reprogramming condition. Mean ± SEM. ****p < 0.0001, one-way ANOVA and Holm-Šídák test in (**H**).

Comparative transcriptomic analysis of ER-T_regs_ and nT_regs_ reveals that ER-T_regs_ exhibit reduced expression of genes linked to T cell quiescence, such as *Lef1*, *Ccr7*, *Bach2*, and *Sell*, and increased expression of T_reg_ effector genes, including *Ctla4*, and *Il10* (69), as well as Th17-associated genes like *Rorc*, *Maf*, *Il23r*, *Ccr6*, and *Il1r1* (70, 71) (Figure 6D). Moreover, ER-T_regs_ express high levels of Th17 markers but not those typical of Th1 or Th2 cells (Figure 6E). GSEA further indicates that ER-T_regs_ upregulate genes involved in effector T_reg_ function, Th17 differentiation, and the IL-23 pathway (Figure 6F), supporting the notion of enhanced Th17 polarization in ER-T_regs_ derived from MOG/CFA-primed T_eff_ cells. Additionally, the observed amplification of SMAD2/3 signaling in ER-T_regs_ suggests that TGF-β in the reprogramming cocktail significantly contributes to their unique gene expression program.

ER-T_regs_ bearing the MOG specific 2D2 TCR express higher levels of c-Maf and Rorγt compared to nT_regs_ with the same 2D2 TCR (72) (Figure 6G), indicating enhanced Th17 polarization in myelin autoantigen-specific ER-T_regs_. To assess the contribution of parental T_eff_ gene expression to this Th17 polarization, we compared Rorγt levels in ER-T_regs_ reprogrammed from in vitro differentiated Th1, Th2, and Th17 cells, as well as in iT_regs_ derived from naïve T cells. ER-T_regs_ reprogrammed from Th17 cells exhibited significantly higher Rorγt expression than those reprogrammed from Th1 cells, Th2 cells, or iT_regs_ (Figure 6H), suggesting that the inheritance of parental Th17 characteristics drives the elevated expression of selective Th17 genes in ER-T_regs_.

### Elevated expression of Th17 genes contributes to ER-T_reg_ fitness and function in EAE

Adoptive transfer of a limited number of ER-T_regs_ (2 × 10⁶ per mouse) into lymphoreplete mice harboring endogenous nT_regs_ significantly ameliorated EAE (Figure 2), demonstrating the superior suppressive capacity of ER-T_regs_ over endogenous nT_regs_. To directly compare their therapeutic efficacy, we transferred MOG/CFA-primed CD4^+^ T_conv_ cells into *Rag1^⁻/⁻^* mice either alone or alongside ER-T_regs_ (derived from MOG/CFA-primed T_eff_ cells) or nT_regs_ (isolated from MOG/CFA-immunized mice). Recipients were immunized with MOG/CFA and treated with PT to induce EAE. ER-T_reg_ co-transfer, but not nT_reg_ co-transfer, effectively suppressed disease progression (Figure 7A). Moreover, ER-T_regs_ exhibited significantly greater accumulation and elevated Ror*γ*t expression in the spinal cord compared to nT_regs_ (Figure 7B–C), suggesting that retained Th17-associated transcriptional programming enhances their tissue fitness.

**Figure 7.**
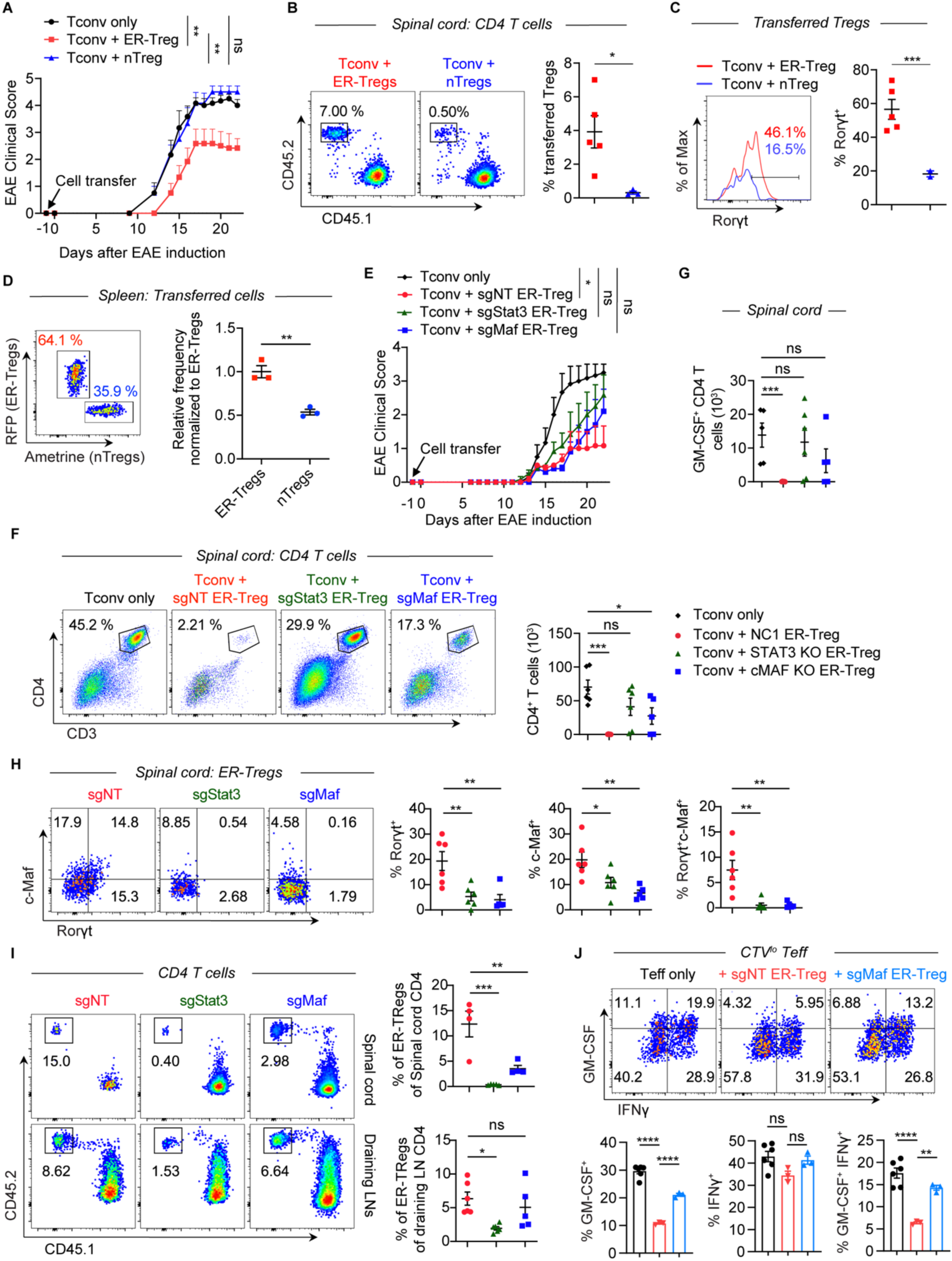
Elevated expression of specific T_eff_ genes contributes to ER-T_reg_ fitness and suppressive function in EAE. (**A** – **C**) EAE was induced in *Rag1^−/−^* mice one day after transfer of MOG/CFA-primed CD45.1^+^ CD4^+^ T_conv_ cells with or without co-transfer of CD45.2^+^ ER-T_regs_ or CD45.2^+^ nT_regs_ isolated from MOG/CFA-primed mice and cultured in the presence of IL-2 for 3 days. Flow cytometry analyses were conducted at 22 dpi. n = 6 per group. (**A**) EAE disease curve. (**B**) Flow cytometry of transferred T_reg_ frequencies. (**C**) Flow cytometry of Rorγt expression in transferred T_regs_. (**D**) Flow cytometry of in vivo competitive fitness between fluorescent reporter transduced 2D2 ER-T_regs_ and nT_regs_, co-transferred at a 1:1 ratio one day prior to CFA/MOG immunization and analyzed 5 days post-immunization. (**E-I**) EAE was induced in *Rag1^−/−^* mice one day after transfer of MOG/CFA-primed CD45.1^+^ CD4^+^ T_conv_ cells with or without co-transfer of CD45.2^+^*Foxp3^Thy1.1^R26^Cas9^*ER-T_regs_ transduced with sgRNA-RV targeting *Stat3* (sgStat3), *Maf* (sgMaf), or a non-targeting sgRNA-RV (sgNT). Flow cytometry analyses were conducted at 22 dpi. n = 6-7 per group. (**E**) EAE disease curve. (**F**) Flow cytometry of the frequencies (left) and numbers (right) of CD4^+^ T cells in the spinal cord. (**G**) Numbers of GM-CSF^+^ CD4^+^ T cells in the spinal cord. (**H**) Flow cytometry of c-Maf and Rorγt expression in draining LN ER-T_regs._ (**I**) Flow cytometry of the frequencies and numbers of transferred ER-T_regs_ in CD4^+^ T cells in the spinal cord (upper) and draining LNs (lower). (**J**) Flow cytometry of IFNγ and GM-CSF expression in CTV^lo^ CD4^+^ T_eff_ cells cocultured for 3 days with APCs and MOG in the presence or absence of sgMaf-RV or sgNT-RV transduced *Foxp3^Thy1.1^R26^Cas9^*ER-T_regs_. Mean ± SEM. *p < 0.05, **p < 0.01, ***p < 0.001, ****p < 0.0001, one-way ANOVA and Holm-Šídák test in (**A**, **E**, **F**, **G**, **H**, **I**, and **J**) and unpaired two-sided t-test in (**B**, **C**, and **D**).

To directly evaluate the role of Th17-related gene expression in ER-T_reg_ fitness, we performed a competitive fitness assay using ER-T_regs_ and nT_regs_ from 2D2 MOG-specific mice. Both cell types were cultured under identical reprogramming conditions, transduced with distinct fluorescent reporters, and co-transferred at a 1:1 ratio into MOG/CFA-immunized mice. ER-T_regs_ outcompeted nT_regs_ in vivo (Figure 7D), confirming that Th17-associated gene signatures enhance their survival and expansion. Collectively, these findings indicate that the inherited Th17-related transcriptional program underpins the enhanced fitness and suppressive efficacy of ER-T_regs_ relative to nT_regs_ in EAE.

To determine whether heightened Th17 polarization underpins ER-T_reg_ functionality in EAE, we used CRISPR/Cas9 to delete the Th17-associated transcription factors STAT3 or c-Maf in ER-Tregs and assessed their suppressive capacity. Co-transfer of control sgNT-transduced *Foxp3^Thy1.1^R26^Cas9^*ER-T_regs_ robustly attenuated EAE progression, whereas ER-T_regs_ lacking STAT3 (sgStat3) or c-Maf (sgMa*f*) failed to suppress disease (Figure 7E). Genetic ablation of *Stat3* or *Maf* impaired the ability of ER-T_regs_ to reduce spinal cord infiltration by total CD4^+^ T cells and GM-CSF^+^CD4^+^ T_eff_ cells (Figure 7F–G), which was accompanied by diminished expression of RORγt and c-Maf in ER-T_regs_ (Figure 7H). Although deletion of *Stat3* or *Maf* reduced the frequency of ER-T_regs_ in the spinal cord (Figure 7I), their absolute numbers remained unchanged or even elevated, respectively (Supplemental Figure 4A), likely due to increased CD4^+^ T cell accumulation in mice receiving knockout ER-T_regs_ (Figure 7E). Notably, Foxp3 expression levels remained consistent across all groups (Supplemental Figure 4B), ruling out gross instability. These findings establish STAT3 and c-Maf as critical drivers of ER-T_reg_ fitness and function, enabling their suppression of neuroinflammation via Th17-associated transcriptional programs.

To determine whether heightened Th17 polarization enhances ER-T_reg_ suppressor function on a per-cell basis, we performed in vitro suppression assays comparing *Maf*-deficient ER-T_regs_ to control ER-T_regs_ transduced with sgNT. Genetic ablation of *Maf* significantly impaired the ability of ER-T_regs_ to suppress GM-CSF^+^CD4^+^ T_eff_ cell responses (Figure 7J), indicating that elevated *Maf* expression is essential for their per-cell suppressive potency. Together, these results demonstrate that Th17-skewed transcriptional program in ER-T_regs_ is critical for their superior tissue fitness and functional efficacy in EAE compared to nT_regs_.

## DISCUSSION

Endogenous T_regs_ acquire their identity through tolerogenic signals that imprint T_reg_-specific transcriptional and epigenetic programs onto naïve precursors. However, whether such programs can be stably established in committed CD4^+^ T_eff_ cells, which retain inflammatory epigenetic memory, remains unknown. To address this, we developed an epigenetic reprogramming strategy to activate core T_reg_ transcriptional circuitry in T_eff_ cells, creating a model system to probe T_reg_ plasticity and therapeutic potential.

*Foxp3* induction in T_eff_ cells is achieved through the synergistic actions of TGF-β, RA, and VC, each contributing distinct mechanistic pathways. TGF-β drives Foxp3 expression via Smad3 binding to the conserved noncoding sequence 1 (CNS1) enhancer of the *Foxp3* locus (73). RA amplifies this process by enhancing TGF-β/Smad3 signaling while concurrently suppressing inflammatory pathways: it downregulates IL-6 and IL-23 receptor expression, neutralizes cytokine-mediated inhibition of *Foxp3* expression, and reduces pro-inflammatory cytokine secretion by T_eff_ cells (74–77). VC stabilizes *Foxp3* expression by promoting TET enzyme-dependent DNA demethylation at TSDRs, including the *Foxp3* locus itself (38, 39, 78). Stable *Foxp3* induction further requires re-stimulation, which sustains transcriptional activation of *Foxp3* and facilitates demethylation of the CNS2 enhancer (49), establishing a self-reinforcing loop to stabilize T_reg_ identity. Notably, our reprogramming approach recapitulates the epigenetic remodeling observed in endogenous T_regs_, inducing robust DNA demethylation at TSDRs of key T_reg_ identity genes such as *Ctla4*, *Il2ra*, and *Ikzf4*. Crucially, this demethylation occurs independently of Foxp3 (16), indicating that TGF-β, RA, and VC cooperatively remodel the epigenome of T_eff_ cells to activate T_reg_ transcriptional programs through both Foxp3-dependent and Foxp3-independent mechanisms. These findings highlight the ability of tolerogenic signals to override inflammatory epigenetic memory in T_eff_ cells, enabling their conversion into functionally stable T_regs_.

Previous studies have established that ectopic Foxp3 expression in conventional CD4^+^ T cells—composed of both naïve and effector T cells—can confer suppressive activity (17–19). However, the use of conventional T cells complicates interpretation of these results. Naïve T cells lack pre-existent inflammatory phenotypes and thus are more amenable to acquiring a suppressive phenotype via Foxp3 overexpression. These Foxp3 overexpressing naïve T cells could mask the resistance of T_eff_ cells, which do have a pre-existent inflammatory phenotype and epigenome, toward acquiring a suppressive phenotype. Whether Foxp3 expression alone suffices to induce authentic T_reg_ phenotypes and suppressive function in T_eff_ cells has thus remained unresolved. Our findings reveal that both stable Foxp3 induction and demethylation of Foxp3-independent T_reg_ identity genes are indispensable for establishing functional T_reg_ programs in T_eff_ cells. Foxp3 critically suppresses IL-17A expression in reprogrammed cells, mirroring its role in pT_regs_ (60), while also enhancing IL-10 production, potentially amplifying their suppressive capacity. Notably, forced Foxp3 expression in T_eff_ cells, without concurrent epigenetic reprogramming, failed to confer suppressor function, underscoring the necessity of Foxp3-independent epigenetic remodeling at T_reg_ identity loci. This aligns with prior work demonstrating that T_reg_-specific DNA demethylation, unattainable through Foxp3 overexpression alone, is essential for T_reg_ functionality (16). Consistent with this, Foxp3-expressing T_eff_ cells exhibited diminished expression of CD25, CTLA-4, and Helios—proteins encoded by genes harboring TSDRs (16)—suggesting that these epigenetic modifications regulate key aspects of T_reg_ gene expression. Collectively, our work highlights a dual requirement for Foxp3 and TSDR-driven epigenetic activation to override T_eff_ cell transcriptional programs and enforce stable T_reg_ identity. Future studies are warranted to dissect the precise contributions of individual TSDRs to T_reg_ transcriptional programs and their therapeutic potential in reprogramming T_eff_ cells to treat autoimmune inflammation.

While MOG-nonspecific ER-T_regs_ alone failed to ameliorate established EAE, bystander suppression mechanisms—potentially mediated by nonspecific ER-T_regs_ in the presence of MOG-specific counterparts—may contribute to disease control (79, 80). Notably, molecular mimicry between myelin antigens and foreign antigens (e.g., Epstein-Barr virus, gut microbiota) has emerged as a key driver of MS/EAE pathogenesis (81–83). Although MOG-specific ER-T_regs_ did not impair OVA-specific vaccine responses, they may suppress immunity against microbial antigens sharing epitopes with autoantigens. These findings highlight the need to rigorously assess the long-term effects of autoantigen-specific ER-T_reg_ therapy on immune homeostasis and infection resilience.

In contrast to ER-T_regs_, adoptive transfer of MOG/CFA-primed nT_regs_ failed to suppress EAE induced by co-transferred MOG/CFA-primed T_conv_ cells in *Rag1^⁻/⁻^* mice. While prior studies reported EAE mitigation using nT_regs_ isolated from naïve mice or recovery-phase mice (84–86), these protocols transferred nT_regs_ into naïve hosts at disease induction—a context lacking pre-existing T_eff_ cell differentiation and inflammation. By contrast, our co-transfer model in *Rag1^⁻/⁻^* mice recapitulates the challenge of suppressing primed T_eff_ cells in an inflammatory milieu. Further, MOG/CFA immunization destabilizes MOG-specific nT_reg_ lineage commitment (7), likely impairing their suppressive capacity upon isolation and transfer. This instability, coupled with intrinsic limitations in effector T_reg_ differentiation, may explain nT_reg_ inefficacy in our model and prior studies (87). Future studies reprogramming destabilized nT_regs_ could clarify their functional potential in EAE suppression.

We observed heightened expression of Rorγt and c-Maf in ER-T_regs_ compared to nT_regs_, even when both cell types expressed the same MOG specific 2D2 TCR. This suggests that the upregulation of Rorγt expression in endogenous nT_regs_, which has been recently shown to be associated with the ability of nT_regs_ to suppress Th17 inflammation in EAE (33), is hindered by the pre-existing gene expression and/or epigenetic landscape in nT_regs_. This phenomenon may serve as a regulatory mechanism preventing endogenous T_regs_ from impeding beneficial anti-pathogen immune responses, while potentially contributing to the development of pathogenic autoimmune inflammation under specific conditions. Further investigations into the molecular mechanisms underlying the constrained or delayed Th17 polarization of nT_regs_ in EAE and its impact on disease progression are warranted and may yield insights into the dysfunction of endogenous T_regs_ in Th17-cell-driven autoimmune diseases.

ER-T_regs_ reprogrammed from MOG/CFA-primed CD4^+^ Teff cells exhibit a unique transcriptional profile, characterized by elevated expression of Th17-associated genes (*Rorc*, *Maf*) compared to nT_regs_. CRISPR-mediated deletion of *Maf* or *Stat3*—key Th17 transcription factors—significantly impaired ER-T_reg_ survival and suppressive function in EAE, highlighting the critical role of Th17-like polarization in their therapeutic efficacy. While prior studies suggest T_regs_ expressing lineage-specific transcription factors (e.g., Th1, Th17) display enhanced suppression of corresponding T_eff_ subsets (23–30), the mechanistic basis remains unclear. Our data suggest that improved cellular fitness, mediated by transcription factor-driven adaptation to inflammatory niches, may underpin this phenomenon.

The heightened Th17 signature in ER-T_regs_ may arise from epigenetic inheritance of parental Th17 cell programs or de novo activation during reprogramming. ER-T_regs_ also displayed amplified SMAD2/3 signaling, indicative of enhanced TGF-β activity, which likely drives *Maf* and *Rorc* expression—both established TGF-β targets in T_regs_ (31, 34, 88, 89). Given the inclusion of TGF-β in the reprogramming cocktail, these findings underscore how lineage-specific epigenetic memory synergizes with extrinsic signals to shape ER-T_reg_ functionality. Future studies should delineate how epigenetic landscapes of distinct T helper subsets influence ER-T_reg_ differentiation and function, particularly in inflammatory contexts.

Our study highlights the superior in vivo fitness of ER-T_regs_ over other T_reg_ subsets, driven by intrinsic properties like their polarization state. CRISPR-mediated ablation of c-Maf impaired ER-T_reg_ suppression of GM-CSF^+^ T_eff_ cells in vitro, suggesting that Th17 polarization enhances their per-cell suppressive potency. While APC interactions were not directly assessed, bulk RNA-seq revealed elevated *Ctla4* expression in ER-T_regs_ (Figure 6D). Given the established role of CTLA-4 in downregulating CD80/CD86 on APCs via transendocytosis (90–93), this upregulation likely contributes to their enhanced suppressive function. Beyond EAE, Th17-derived ER-T_regs_—combining stable Foxp3 expression, robust tissue persistence, and potent suppression—hold promise for other Th17-driven diseases such as inflammatory bowel disease. More broadly, our epigenetic reprogramming strategy could be applied to other helper subsets; for example, converting autoreactive Th1 cells into stable T_regs_ may offer a targeted approach to mitigate pancreatic β-cell autoimmunity in type 1 diabetes.

Our study establishes that coordinated demethylation of *Foxp3* and Foxp3-independent T_reg_ identity genes enables the conversion of committed CD4^+^ T_eff_ cells into functional T_regs_ with bona fide transcriptional and suppressive programs. The resulting ER-T_regs_ outperform endogenous nT_regs_, iT_regs_, and Foxp3-overexpressing T_eff_ cells in suppressing established autoimmune inflammation, underscoring their therapeutic potential. This enhanced efficacy arises from dual mechanisms: (1) inherited autoantigen specificity, which promotes antigen specific and tissue-localized suppression, and (2) retention of parental T_eff_ transcriptional programs, which bolsters fitness in inflammatory niches. These insights advance our understanding of T_reg_ epigenetic regulation while offering a blueprint for engineering antigen-specific T_reg_ therapies with tailored functionality to treat autoimmune diseases.

## METHODS

### Sex as a biological variable

Though women are more susceptible to developing MS than men by a ratio of approximately 3:1, men that develop MS exhibit greater cognitive impairment and more rapid disability progression than women. Our study examined both male and female mice and similar findings are reported for both sexes.

### Mice

Animals were housed at the Tufts University School of Medicine (TUSM) animal facility under specific pathogen-free conditions according to institutional guidelines. All studies were performed under protocol B2022-85 and approved by Tufts Institutional Animal Care and Use Committee. All mouse strains used were on the C57BL/6 genetic background. *CD45.1* (#002014), *Rosa26^Cas9-eGFP^* (#026179), *2D2* (#006912), and *Rag1^-/-^* (#002216) mice were purchased from The Jackson Laboratory. *Foxp3^Thy1.1^* mice were a gift from Dr. Y. Zheng. The above mouse strains were bred in-house at TUSM to produce the *Foxp3^Thy1.1^Rosa26^Cas9^*, *Foxp3^Thy1.1^CD45.1^+/+^*, and *2D2^+/-^ Foxp3^Thy1.1^CD45.1^+/-^* mouse strains. Male and female mice used were at least 6 weeks old and had no prior exposure to drugs or experimentation.

### Antibodies and reagents

Flow cytometry antibodies anti-CD3 (2C11), anti-CD4 (RM4-5), anti-Thy1.1 (HIS51), anti-CD44 (IM7), anti-CD62L (MEL-14), anti-CD45.1 (A20), anti-CD45.2 (104), anti-IFNγ (XMG1.2), anti-GMCSF (MP1-22E9), anti-B220 (RA3-6B2), anti-GL7 (GL7), anti-CD138 (281–2), anti-CXCR5 (L138D7), and anti-NGFR (ME20.4) were purchased from BioLegend. Anti-IL-17 (eBio17B7), anti-Foxp3 (FJK-16s), anti-Rorγt (B2D), and anti-c-MAF (sym0F1) were purchased from eBioscience. Anti-CD95 (Jo2) was purchased from BD Biosciences.

Neutralizing antibodies toward IFNγ (XMG1.2), IL-4 (11B11), and IL-12 (C17.8) were purchased from BioXcell. Human IL-2 and IL-7 were purchased from PeproTech. Mouse TGFb, IL-6, IL-23, and IL-1b were purchased from R&D systems. Retinoic acid was purchased from Sigma-Aldrich. Vitamin C was purchased from Fisher Scientific. CD3/CD28 Dynabeads were purchased from Thermo Fisher Scientific. Incomplete Fruend’s Adjuvant was purchased from Thermo Fisher Scientific. Myelin oligodendrocyte glycoprotein (amino acids 35-55) was purchased from GeneMed Synthesis. Heat killed *Mycobacterium tuberculosis* strain H37 Ra was purchased from BD Biosciences.

### ER-T_reg_ generation

CD4^+^CD44^hi^ T_eff_ were sort purified from donor mice that were immunized with CFA/MOG 7 days prior. Sorted cells were rested in T cell growth medium (RPMI 1640 supplemented with 2mM GlutaMAX, 10mM HEPES, 100 U/mL penicillin/streptomycin, 1 mM sodium pyruvate, 5% fetal calf serum) for 4 days in the presence of 2 ng/mL IL-7 and 10 ug/mL neutralizing antibodies against IFNγ, IL-4, and IL-12. Cells were then stimulated with CD3/CD28 Dynabeads for 4 days in T cell growth medium supplemented with 1,000 U/mL IL-2, 5 ng/mL TGFb, 100 ug/mL vitamin C, 10 nM retinoic acid, and 10 ug/mL cytokine neutralizing antibodies (primary reprogramming cocktail). CD4^+^Thy1.1^+^ cells were sort purified and restimulated with Dynabeads in T cell growth medium supplemented with 1,000 U/mL IL-2, 2 ng/mL TGFb, 10 ug/mL vitamin C, and 10 ug/mL cytokine neutralizing antibodies (secondary reprogramming cocktail). For experiments using transduced ER-T_regs_, CD4^+^Thy1.1^+^Reporter^+^ cells were sort purified and restimulated.

### Retroviral vectors

MG2A and MG2N were generated by modifying MSCV-P2GM-FF (plasmid no. 19750, Addgene). Mouse *Foxp3, Maf, Rorc,* and *Stat3* single guide RNAs (sgRNA) were cloned into BbsI-digested MG2N or MG2A. MIGR-mFoxp3 was a gift from D. Littman (plasmid no. 24067, Addgene). The guide sequences are nontargeting (NT) (5’-GCACTACCAGAGCTAACTCA-3’), *Foxp3* (5’-GTTCCTGGGTGTACCCGAGCG-3’), *Maf* (5’-GCCCGCAGCAGCTCAACCCGG-3’), *Rorc* (5’-GTCATCTGGGATCCACTACG-3’), and *Stat3* (5’-GAGATTATGAAACACCAACG-3’).

### Production of retrovirus

Retrovirus was produced by transfecting HEK293T cells (ATCC) 2 days prior to transduction using the pCL-Eco packaging vector and Fugene HD transfection reagent (Promega). Medium was replaced with half the transfection volume 1 day before transduction.

### Retroviral transduction

Rested CD4^+^ T_eff_ cells were stimulated for 1 day with CD3/CD28 Dynabeads in the presence of the primary reprogramming cocktail. Cells were then transduced by spin infection with viral supernatant supplemented with the primary reprogramming cocktail and 4 ug/mL polybrene. Spin infection was performed in a Sorvall Legend X1R centrifuge for 90 minutes at 2,800 RPM and 37C.

### Bisulfite sequencing

Genomic DNA was isolated using the GeneJet Genomic DNA Purification Kit (Thermo Fisher Scientific) and then bisulfute converted using EpiTect Bisufite Conversion Kit (Qiagen). Converted DNA was amplified with Q5U polymerase (New England Biolabs) and gel purified after agarose gel electrophoresis. Purified PCR product was cloned into pJET1.2 (Thermo Fisher Scientific) for Sanger sequencing. The bisulfite amplification primers are CNS2 Forward (5’ - TGGGTTTTTTTGGTATTTAAGAAAG-3’), CNS2 Reverse (5’-AACCAACCAACTTCCTACACTATCTAT-3’), CTLA4 Forward (5’-TGGTGTTGGTTAGTAGTTATGGTGT-3’), CTLA4 Reverse (5’-AAATTCCACCTTACAAAAATACAATC-3’), IL2ra Forward (5’-TTTTAGAGTTAGAAGATAGAAGGTATGGAA-3’), IL2ra Reverse (5’-TCCCAATACTTAACAAAACCACATAT-3’), Ikzf4 Forward (5’-AGGATGGTTTTTATTGAAGGTGAT-3’), Ikzf4 Reverse (5’-ATACACACCAAACAAACACTACACC-3’).

### EAE induction

EAE was induced by subcutaneous injection of 50 uL of an emulsion containing 50 ug MOG_35-55_ and 250 ug *M. tuberculosis* strain H37 Ra in Incomplete Fruend’s Adjuvant into each hind flank. Mice also received an intraperitoneal injection of 200 ng pertussis toxin in 200 uL PBS on days 0 and 2 after immunization. Clinical signs of EAE were assessed by the following criteria: 0, no signs of disease; 1, loss of tail tone; 2, hind limb paresis; 3, hind limb paralysis; 4, tetraplegia; 5, moribund or dead. Mice with a score greater than 4 were euthanized and carried with a score of 5 for the duration of the experiment.

### Cell transfer

For all preventive EAE experiments in *Rag1^-/-^* mice, cells were transferred intravenously one day prior to disease initiation. Mice received 50,000 T_regs_ and 100,000 CD4^+^ T_conv_ from CD45 congenically distinct *Foxp3^Thy1.1^* donor mice. CD4^+^ T_conv_ were procured from mice which were immunized with CFA/MOG 7 days prior to transfer using the mouse CD4^+^ T Cell Isolation Kit (Miltenyi Biotec) followed by T_reg_ depletion using anti-Thy1.1 PE (HIS51) and anti-PE nanobeads (BioLegend). For preventive EAE experiments in lymphoreplete mice, 2 x 10^6^ T_regs_ from CD45 congenically distinct *Foxp3^Thy1.1^* donor mice were transferred intravenously 1 day prior to disease initiation. For therapeutic EAE experiments in lymphoreplete mice, 2 x 10^6^ polyclonal or 0.5 x 10^6^ MOG-specific CD45 congenically distinct T_regs_ were transferred intravenously at first evidence of disease (tail paralysis, typically day 11 after disease initiation). For in vivo fitness experiments in lymphoreplete mice, 0.5 x 10^6^ 2D2 TCR transgenic CD45 congenically distinct T_regs_ were transferred intravenously one day prior to immunization with MOG/CFA.

### Cell isolation

For lymph node and spleen, tissues were mechanically dissociated using the back of a syringe plunger and filtered through a 70 micron nylon mesh. Spinal cords were harvested by perfusing mice with PBS and collecting the tissue. Spinal cords were then cut into small pieces and enzymatically digested with 1 mg/mL collagenase D and 0.1 mg/mL DNase (Sigma Aldrich) in T cell growth medium for 30 minutes at 37C with shaking at 1,500 RPM. Tissue digests were filtered through a 40 micron nylon mesh and remaining tissue was mechanically dissociated with the back of a syringe plunger. Dissociated spinal cord cells were then placed into a 30%-37%-70% isotonic Percoll (Cytiva) gradient and centrifuged for 30 minutes at room temperature at 800xg to enrich the infiltrating mononuclear cells.

### Flow cytometry

For surface staining, cells were stained in FACS buffer (PBS, 0.5% BSA, 1mM EDTA) for 15 minutes at 4C, washed, and analyzed. For intracellular cytokine staining, cells were incubated at 37C in the presence of 50 ng/mL PMA and 500 ng/mL ionomycin for one hour. GolgiStop (BD Biosciences) was added, and the cells were incubated at 37C for an additional 3 hours.

Stimulated cells were surface stained, fixed and permeabilized with Foxp3/Transcription Factor Staining Kit (Tonbo Biosciences) according to manufacturer instructions, and stained for cytokines in permeabilization buffer. Washed cells were stored in FACS buffer until analysis.

### In vitro suppression assays

CD45 congenically distinct CD4^+^CD44^hi^ T_eff_ cells were sorted from donor mice immunized 7-10 days prior and labeled with CellTrace Violet (Invitrogen). Labeled cells were co-cultured with T cell depleted APCs and T_regs_ in the presence of 10 ug/mL MOG_35-55_. Dilution of CellTrace Violet and cytokine expression were measured 4 days later.

### RNA sequencing

CD45 congenically distinct ER-T_regs_ were transferred to lymphoreplete mice, and the mice were immunized with CFA/MOG. After 7 days, ER-T_regs_, nT_regs_, and CD4^+^ T_eff_ cells were sorted from the draining lymph nodes directly into TRIzol (Qiagen). RNA was isolated using phenol-chloroform extraction. Uniquely indexed libraries were pooled in equimolar ratios and sequenced on a single Illumina NextSeq500 run with single-end 75-bp reads by the Tufts University Genomics Core Facilities.

### RNA-seq analysis

Sequence reads were aligned with the mm39 reference genome assembly and gene counts were quantified with FeatureCounts. Differential expression analysis was performed with DESeq2. Gene-set enrichment analyses were performed with GSEAPreranked, in which genes ranked according to their fold changes were compared with the following MSigDB signature collections: GSE7852_T_reg__VS_T_conv__DN gene set, GSE7852_T_reg__VS_T_conv__UP gene set, WP_TH17_CELL_DIFFERENTIATION_PATHWAY, PID_IL23_PATHWAY, PID_SMAD2_3NUCLEAR_PATHWAY, as well as a CNS2_Dependent_Effector_T_reg_ gene set generated from GSE57272.

### Statistics

Except for RNA-Seq analysis, statistical significance was determined using GraphPad Prism 10.0 (GraphPad Software). For comparisons of a single variable between 2 groups, significance was determined using unpaired t tests. For comparisons of multiple groups where variance did not significantly differ across groups, 1- or 2-way ANOVA with Šidák’s (for comparisons between preselected pairs) multiple-comparison corrections was used. EAE disease scores were analyzed with 1-way ANOVA of area under the curve of clinical scores. *P* values below 0.05 were considered statistically significant and are shown by the exact number or by asterisks in the figures.

### Study approval

All studies were performed under protocol B2022-85 and approved by Tufts Institutional Animal Care and Use Committee.

### Data and materials availability

All data are available in the main text or the Supplemental materials. Individual data point values can be found in the “Supporting Data Values” document associated with this manuscript. RNA sequencing data can be accessed using GEO Accession Number GSE303685. All materials used or generated in this study are available to researchers following appropriate standard material transfer agreements.

## Author contributions

Conceptualization: X.L. and T.R.C.

Methodology: X.L. and T.R.C.

Formal Analysis: T.R.C.

Investigation: T.R.C., J.J.C., H.I.M., M.N., J.L.L., J.H.K., and X.L.

Writing: X.L. and T.R.C.

Visualization: T.R.C.

Supervision: X.L.

Project administration: X.L.

Funding acquisition: X.L.

## Acknowledgments

We thank S. Kwok and A. Parmelee for assistance in flow cytometry, A. Tai and I. Grinvald for assistance in RNA-seq experiments, Y. Zheng and A. Rudensky for sharing the Foxp3Thy1.1 mouse strain for this study, and P. Alcaide and R. Isberg for helpful discussions. T.R.C. was supported by the NIH grant AI167245. J.J.C. was supported by the Tufts University Graduate School of Biomedical Sciences Dean’s Fellowship. X.L. was supported by the Ellison Foundation and the NIH grant AI167245. This work was also supported by the NIH-funded Tufts University Core Facility (grant nos.: 1S10OD016196-01 and S10OD032201).

## References

1. Josefowicz SZ, Lu LF, and Rudensky AY. Regulatory T cells: mechanisms of differentiation and function. Annual review of immunology. 2012;30:531–64.

2. Sakaguchi S, Yamaguchi T, Nomura T, and Ono M. Regulatory T cells and immune tolerance. Cell. 2008;133(5):775–87.

3. Viglietta V, Baecher-Allan C, Weiner HL, and Hafler DA. Loss of functional suppression by CD4+CD25+ regulatory T cells in patients with multiple sclerosis. J Exp Med. 2004;199(7):971–9.

4. Haas J, Hug A, Viehover A, Fritzsching B, Falk CS, Filser A, et al. Reduced suppressive effect of CD4+CD25high regulatory T cells on the T cell immune response against myelin oligodendrocyte glycoprotein in patients with multiple sclerosis. Eur J Immunol. 2005;35(11):3343–52.

5. Raffin C, Vo LT, and Bluestone JA. Treg cell-based therapies: challenges and perspectives. Nat Rev Immunol. 2020;20(3):158–72.

6. Dominguez-Villar M, and Hafler DA. Regulatory T cells in autoimmune disease. Nat Immunol. 2018;19(7):665–73.

7. Bailey-Bucktrout SL, Martinez-Llordella M, Zhou X, Anthony B, Rosenthal W, Luche H, et al. Self-antigen-driven activation induces instability of regulatory T cells during an inflammatory autoimmune response. Immunity. 2013;39(5):949–62.

8. Zhou X, Bailey-Bucktrout SL, Jeker LT, Penaranda C, Martinez-Llordella M, Ashby M, et al. Instability of the transcription factor Foxp3 leads to the generation of pathogenic memory T cells in vivo. Nat Immunol. 2009;10(9):1000–7.

9. Wan YY, and Flavell RA. Regulatory T-cell functions are subverted and converted owing to attenuated Foxp3 expression. Nature. 2007;445(7129):766–70.

10. Huan J, Culbertson N, Spencer L, Bartholomew R, Burrows GG, Chou YK, et al. Decreased FOXP3 levels in multiple sclerosis patients. J Neurosci Res. 2005;81(1):45–52.

11. Joudi AM, Reyes Flores CP, and Singer BD. Epigenetic Control of Regulatory T Cell Stability and Function: Implications for Translation. Front Immunol. 2022;13:861607.

12. Sakaguchi S, Kawakami R, and Mikami N. Treg-based immunotherapy for antigen-specific immune suppression and stable tolerance induction: a perspective. Immunother Adv. 2023;3(1):ltad007.

13. Mikami N, Kawakami R, Chen KY, Sugimoto A, Ohkura N, and Sakaguchi S. Epigenetic conversion of conventional T cells into regulatory T cells by CD28 signal deprivation. Proc Natl Acad Sci U S A. 2020;117(22):12258–68.

14. Akamatsu M, Mikami N, Ohkura N, Kawakami R, Kitagawa Y, Sugimoto A, et al. Conversion of antigen-specific effector/memory T cells into Foxp3-expressing Treg cells by inhibition of CDK8/19. Sci Immunol. 2019;4(40).

15. Williams LM, and Rudensky AY. Maintenance of the Foxp3-dependent developmental program in mature regulatory T cells requires continued expression of Foxp3. Nat Immunol. 2007;8(3):277–84.

16. Ohkura N, Hamaguchi M, Morikawa H, Sugimura K, Tanaka A, Ito Y, et al. T cell receptor stimulation-induced epigenetic changes and Foxp3 expression are independent and complementary events required for Treg cell development. Immunity. 2012;37(5):785–99.

17. Fontenot JD, Gavin MA, and Rudensky AY. Foxp3 programs the development and function of CD4+CD25+ regulatory T cells. Nat Immunol. 2003;4(4):330–6.

18. Hori S, Nomura T, and Sakaguchi S. Control of regulatory T cell development by the transcription factor Foxp3. Science. 2003;299(5609):1057–61.

19. Khattri R, Cox T, Yasayko SA, and Ramsdell F. An essential role for Scurfin in CD4+CD25+ T regulatory cells. Nat Immunol. 2003;4(4):337–42.

20. Li X, and Zheng Y. Regulatory T cell identity: formation and maintenance. Trends in immunology. 2015;36(6):344–53.

21. Dikiy S, and Rudensky AY. Principles of regulatory T cell function. Immunity. 2023;56(2):240–55.

22. Zemmour D, Zilionis R, Kiner E, Klein AM, Mathis D, and Benoist C. Single-cell gene expression reveals a landscape of regulatory T cell phenotypes shaped by the TCR. Nat Immunol. 2018;19(3):291–301.

23. Zheng Y, Chaudhry A, Kas A, deRoos P, Kim JM, Chu TT, et al. Regulatory T-cell suppressor program co-opts transcription factor IRF4 to control T(H)2 responses. Nature. 2009;458(7236):351‒6.

24. Cretney E, Xin A, Shi W, Minnich M, Masson F, Miasari M, et al. The transcription factors Blimp-1 and IRF4 jointly control the differentiation and function of effector regulatory T cells. Nat Immunol. 2011;12(4):304–11.

25. Koch MA, Tucker-Heard G, Perdue NR, Killebrew JR, Urdahl KB, and Campbell DJ. The transcription factor T-bet controls regulatory T cell homeostasis and function during type 1 inflammation. Nat Immunol. 2009;10(6):595–602.

26. Levine AG, Mendoza A, Hemmers S, Moltedo B, Niec RE, Schizas M, et al. Stability and function of regulatory T cells expressing the transcription factor T-bet. Nature. 2017;546(7658):421–5.

27. Wang Y, Su MA, and Wan YY. An essential role of the transcription factor GATA-3 for the function of regulatory T cells. Immunity. 2011;35(3):337–48.

28. Wohlfert EA, Grainger JR, Bouladoux N, Konkel JE, Oldenhove G, Ribeiro CH, et al. GATA3 controls Foxp3(+) regulatory T cell fate during inflammation in mice. J Clin Invest. 2011;121(11):4503–15.

29. Ohnmacht C, Park JH, Cording S, Wing JB, Atarashi K, Obata Y, et al. MUCOSAL IMMUNOLOGY. The microbiota regulates type 2 immunity through RORgammat(+) T cells. Science. 2015;349(6251):989–93.

30. Sefik E, Geva-Zatorsky N, Oh S, Konnikova L, Zemmour D, McGuire AM, et al. MUCOSAL IMMUNOLOGY. Individual intestinal symbionts induce a distinct population of RORgamma(+) regulatory T cells. Science. 2015;349(6251):993–7.

31. Xu M, Pokrovskii M, Ding Y, Yi R, Au C, Harrison OJ, et al. c-MAF-dependent regulatory T cells mediate immunological tolerance to a gut pathobiont. Nature. 2018;554(7692):373–7.

32. Neumann C, Blume J, Roy U, Teh PP, Vasanthakumar A, Beller A, et al. c-Maf-dependent T(reg) cell control of intestinal T(H)17 cells and IgA establishes host-microbiota homeostasis. Nat Immunol. 2019;20(4):471–81.

33. Kim BS, Lu H, Ichiyama K, Chen X, Zhang YB, Mistry NA, et al. Generation of RORgammat(+) Antigen-Specific T Regulatory 17 Cells from Foxp3(+) Precursors in Autoimmunity. Cell Rep. 2017;21(1):195–207.

34. Wheaton JD, Yeh CH, and Ciofani M. Cutting Edge: c-Maf Is Required for Regulatory T Cells To Adopt RORgammat(+) and Follicular Phenotypes. J Immunol. 2017;199(12):3931–6.

35. Wang L, Liu Y, Beier UH, Han R, Bhatti TR, Akimova T, et al. Foxp3+ T-regulatory cells require DNA methyltransferase 1 expression to prevent development of lethal autoimmunity. Blood. 2013;121(18):3631–9.

36. Helmin KA, Morales-Nebreda L, Torres Acosta MA, Anekalla KR, Chen SY, Abdala-Valencia H, et al. Maintenance DNA methylation is essential for regulatory T cell development and stability of suppressive function. J Clin Invest. 2020;130(12):6571–87.

37. Yang R, Qu C, Zhou Y, Konkel JE, Shi S, Liu Y, et al. Hydrogen Sulfide Promotes Tet1- and Tet2-Mediated Foxp3 Demethylation to Drive Regulatory T Cell Differentiation and Maintain Immune Homeostasis. Immunity. 2015;43(2):251–63.

38. Yue X, Trifari S, Aijo T, Tsagaratou A, Pastor WA, Zepeda-Martinez JA, et al. Control of Foxp3 stability through modulation of TET activity. J Exp Med. 2016;213(3):377–97.

39. Sasidharan Nair V, Song MH, and Oh KI. Vitamin C Facilitates Demethylation of the Foxp3 Enhancer in a Tet-Dependent Manner. J Immunol. 2016;196(5):2119–31.

40. Morikawa H, Ohkura N, Vandenbon A, Itoh M, Nagao-Sato S, Kawaji H, et al. Differential roles of epigenetic changes and Foxp3 expression in regulatory T cell-specific transcriptional regulation. Proc Natl Acad Sci U S A. 2014;111(14):5289–94.

41. Floess S, Freyer J, Siewert C, Baron U, Olek S, Polansky J, et al. Epigenetic control of the foxp3 locus in regulatory T cells. PLoS Biol. 2007;5(2):e38.

42. Polansky JK, Kretschmer K, Freyer J, Floess S, Garbe A, Baron U, et al. DNA methylation controls Foxp3 gene expression. Eur J Immunol. 2008;38(6):1654–63.

43. Singer BD, Mock JR, Aggarwal NR, Garibaldi BT, Sidhaye VK, Florez MA, et al. Regulatory T cell DNA methyltransferase inhibition accelerates resolution of lung inflammation. Am J Respir Cell Mol Biol. 2015;52(5):641–52.

44. Li X, Liang Y, LeBlanc M, Benner C, and Zheng Y. Function of a Foxp3 cis-element in protecting regulatory T cell identity. Cell. 2014;158(4):734–48.

45. Feng Y, Arvey A, Chinen T, van der Veeken J, Gasteiger G, and Rudensky AY. Control of the inheritance of regulatory T cell identity by a cis element in the Foxp3 locus. Cell. 2014;158(4):749–63.

46. Dardalhon V, Awasthi A, Kwon H, Galileos G, Gao W, Sobel RA, et al. IL-4 inhibits TGF-beta-induced Foxp3+ T cells and, together with TGF-beta, generates IL-9+ IL-10+ Foxp3(-) effector T cells. Nat Immunol. 2008;9(12):1347–55.

47. Korn T, Mitsdoerffer M, Croxford AL, Awasthi A, Dardalhon VA, Galileos G, et al. IL-6 controls Th17 immunity in vivo by inhibiting the conversion of conventional T cells into Foxp3+ regulatory T cells. Proc Natl Acad Sci U S A. 2008;105(47):18460–5.

48. Wei J, Duramad O, Perng OA, Reiner SL, Liu YJ, and Qin FX. Antagonistic nature of T helper 1/2 developmental programs in opposing peripheral induction of Foxp3+ regulatory T cells. Proc Natl Acad Sci U S A. 2007;104(46):18169–74.

49. Cameron J, Martino P, Nguyen L, and Li X. Cutting Edge: CRISPR-Based Transcriptional Regulators Reveal Transcription-Dependent Establishment of Epigenetic Memory of Foxp3 in Regulatory T Cells. J Immunol. 2020;205(11):2953–8.

50. Liston A, Nutsch KM, Farr AG, Lund JM, Rasmussen JP, Koni PA, et al. Differentiation of regulatory Foxp3+ T cells in the thymic cortex. Proc Natl Acad Sci U S A. 2008;105(33):11903–8.

51. Chen W, Jin W, Hardegen N, Lei KJ, Li L, Marinos N, et al. Conversion of peripheral CD4+CD25-naive T cells to CD4+CD25+ regulatory T cells by TGF-beta induction of transcription factor Foxp3. J Exp Med. 2003;198(12):1875–86.

52. Fu S, Zhang N, Yopp AC, Chen D, Mao M, Chen D, et al. TGF-beta induces Foxp3 + T-regulatory cells from CD4 + CD25 - precursors. American journal of transplantation : oIicial journal of the American Society of Transplantation and the American Society of Transplant Surgeons. 2004;4(10):1614–27.

53. Coombes JL, Siddiqui KR, Arancibia-Carcamo CV, Hall J, Sun CM, Belkaid Y, et al. A functionally specialized population of mucosal CD103+ DCs induces Foxp3+ regulatory T cells via a TGF-beta and retinoic acid-dependent mechanism. J Exp Med. 2007;204(8):1757–64.

54. Sun CM, Hall JA, Blank RB, Bouladoux N, Oukka M, Mora JR, et al. Small intestine lamina propria dendritic cells promote de novo generation of Foxp3 T reg cells via retinoic acid. J Exp Med. 2007;204(8):1775–85.

55. Benson MJ, Pino-Lagos K, Rosemblatt M, and Noelle RJ. All-trans retinoic acid mediates enhanced T reg cell growth, differentiation, and gut homing in the face of high levels of co-stimulation. J Exp Med. 2007;204(8):1765–74.

56. Mucida D, Park Y, Kim G, Turovskaya O, Scott I, Kronenberg M, et al. Reciprocal TH17 and regulatory T cell differentiation mediated by retinoic acid. Science. 2007;317(5835):256–60.

57. Ponomarev ED, Shriver LP, Maresz K, Pedras-Vasconcelos J, Verthelyi D, and Dittel BN. GM-CSF production by autoreactive T cells is required for the activation of microglial cells and the onset of experimental autoimmune encephalomyelitis. J Immunol. 2007;178(1):39–48.

58. McQualter JL, Darwiche R, Ewing C, Onuki M, Kay TW, Hamilton JA, et al. Granulocyte macrophage colony-stimulating factor: a new putative therapeutic target in multiple sclerosis. J Exp Med. 2001;194(7):873–82.

59. Croxford AL, Lanzinger M, Hartmann FJ, Schreiner B, Mair F, Pelczar P, et al. The Cytokine GM-CSF Drives the Inflammatory Signature of CCR2+ Monocytes and Licenses Autoimmunity. Immunity. 2015;43(3):502–14.

60. van der Veeken J, Campbell C, Pritykin Y, Schizas M, Verter J, Hu W, et al. Genetic tracing reveals transcription factor Foxp3-dependent and Foxp3-independent functionality of peripherally induced Treg cells. Immunity. 2022;55(7):1173–84 e7.

61. Wing K, Onishi Y, Prieto-Martin P, Yamaguchi T, Miyara M, Fehervari Z, et al. CTLA-4 control over Foxp3+ regulatory T cell function. Science. 2008;322(5899):271–5.

62. Chinen T, Kannan AK, Levine AG, Fan X, Klein U, Zheng Y, et al. An essential role for the IL-2 receptor in Treg cell function. Nat Immunol. 2016;17(11):1322–33.

63. Kim HJ, Barnitz RA, Kreslavsky T, Brown FD, Moffett H, Lemieux ME, et al. Stable inhibitory activity of regulatory T cells requires the transcription factor Helios. Science. 2015;350(6258):334–9.

64. Sebastian M, Lopez-Ocasio M, Metidji A, Rieder SA, Shevach EM, and Thornton AM. Helios Controls a Limited Subset of Regulatory T Cell Functions. J Immunol. 2016;196(1):144–55.

65. Fontenot JD, Rasmussen JP, Gavin MA, and Rudensky AY. A function for interleukin 2 in Foxp3-expressing regulatory T cells. Nat Immunol. 2005;6(11):1142–51.

66. D’Cruz LM, and Klein L. Development and function of agonist-induced CD25+Foxp3+ regulatory T cells in the absence of interleukin 2 signaling. Nat Immunol. 2005;6(11):1152–9.

67. Levine AG, Arvey A, Jin W, and Rudensky AY. Continuous requirement for the TCR in regulatory T cell function. Nat Immunol. 2014;15(11):1070–8.

68. Vahl JC, Drees C, Heger K, Heink S, Fischer JC, Nedjic J, et al. Continuous T cell receptor signals maintain a functional regulatory T cell pool. Immunity. 2014;41(5):722–36.

69. Miragaia RJ, Gomes T, Chomka A, Jardine L, Riedel A, Hegazy AN, et al. Single-Cell Transcriptomics of Regulatory T Cells Reveals Trajectories of Tissue Adaptation. Immunity. 2019;50(2):493–504 e7.

70. Lee Y, Awasthi A, Yosef N, Quintana FJ, Xiao S, Peters A, et al. Induction and molecular signature of pathogenic TH17 cells. Nat Immunol. 2012;13(10):991–9.

71. Ciofani M, Madar A, Galan C, Sellars M, Mace K, Pauli F, et al. A validated regulatory network for Th17 cell specification. Cell. 2012;151(2):289–303.

72. Bettelli E, Pagany M, Weiner HL, Linington C, Sobel RA, and Kuchroo VK. Myelin oligodendrocyte glycoprotein-specific T cell receptor transgenic mice develop spontaneous autoimmune optic neuritis. J Exp Med. 2003;197(9):1073–81.

73. Tone Y, Furuuchi K, Kojima Y, Tykocinski ML, Greene MI, and Tone M. Smad3 and NFAT cooperate to induce Foxp3 expression through its enhancer. Nat Immunol. 2008;9(2):194–202.

74. Xiao S, Jin H, Korn T, Liu SM, Oukka M, Lim B, et al. Retinoic acid increases Foxp3+ regulatory T cells and inhibits development of Th17 cells by enhancing TGF-beta-driven Smad3 signaling and inhibiting IL-6 and IL-23 receptor expression. J Immunol. 2008;181(4):2277–84.

75. Mucida D, Pino-Lagos K, Kim G, Nowak E, Benson MJ, Kronenberg M, et al. Retinoic acid can directly promote TGF-beta-mediated Foxp3(+) Treg cell conversion of naive T cells. Immunity. 2009;30(4):471–2; author reply 2-3.

76. Takaki H, Ichiyama K, Koga K, Chinen T, Takaesu G, Sugiyama Y, et al. STAT6 Inhibits TGF-beta1-mediated Foxp3 induction through direct binding to the Foxp3 promoter, which is reverted by retinoic acid receptor. J Biol Chem. 2008;283(22):14955–62.

77. Nolting J, Daniel C, Reuter S, Stuelten C, Li P, Sucov H, et al. Retinoic acid can enhance conversion of naive into regulatory T cells independently of secreted cytokines. J Exp Med. 2009;206(10):2131–9.

78. Blaschke K, Ebata KT, Karimi MM, Zepeda-Martinez JA, Goyal P, Mahapatra S, et al. Vitamin C induces Tet-dependent DNA demethylation and a blastocyst-like state in ES cells. Nature. 2013;500(7461):222–6.

79. Krienke C, Kolb L, Diken E, Streuber M, Kirchhoff S, Bukur T, et al. A noninflammatory mRNA vaccine for treatment of experimental autoimmune encephalomyelitis. Science. 2021;371(6525):145–53.

80. Kenison JE, Jhaveri A, Li Z, Khadse N, Tjon E, Tezza S, et al. Tolerogenic nanoparticles suppress central nervous system inflammation. Proc Natl Acad Sci U S A. 2020;117(50):32017–28.

81. Lanz TV, Brewer RC, Ho PP, Moon JS, Jude KM, Fernandez D, et al. Clonally expanded B cells in multiple sclerosis bind EBV EBNA1 and GlialCAM. Nature. 2022;603(7900):321–7.

82. Tengvall K, Huang J, Hellstrom C, Kammer P, Bistrom M, Ayoglu B, et al. Molecular mimicry between Anoctamin 2 and Epstein-Barr virus nuclear antigen 1 associates with multiple sclerosis risk. Proc Natl Acad Sci U S A. 2019;116(34):16955–60.

83. Bjornevik K, Cortese M, Healy BC, Kuhle J, Mina MJ, Leng Y, et al. Longitudinal analysis reveals high prevalence of Epstein-Barr virus associated with multiple sclerosis. Science. 2022;375(6578):296–301.

84. Kohm AP, Carpentier PA, Anger HA, and Miller SD. Cutting edge: CD4+CD25+ regulatory T cells suppress antigen-specific autoreactive immune responses and central nervous system inflammation during active experimental autoimmune encephalomyelitis. J Immunol. 2002;169(9):4712–6.

85. Zhang X, Koldzic DN, Izikson L, Reddy J, Nazareno RF, Sakaguchi S, et al. IL-10 is involved in the suppression of experimental autoimmune encephalomyelitis by CD25+CD4+ regulatory T cells. Int Immunol. 2004;16(2):249–56.

86. McGeachy MJ, Stephens LA, and Anderton SM. Natural recovery and protection from autoimmune encephalomyelitis: contribution of CD4+CD25+ regulatory cells within the central nervous system. J Immunol. 2005;175(5):3025–32.

87. Korn T, Reddy J, Gao W, Bettelli E, Awasthi A, Petersen TR, et al. Myelin-specific regulatory T cells accumulate in the CNS but fail to control autoimmune inflammation. Nature medicine. 2007;13(4):423–31.

88. Konkel JE, Zhang D, Zanvit P, Chia C, Zangarle-Murray T, Jin W, et al. Transforming Growth Factor-beta Signaling in Regulatory T Cells Controls T Helper-17 Cells and Tissue-Specific Immune Responses. Immunity. 2017;46(4):660–74.

89. Chang D, Xing Q, Su Y, Zhao X, Xu W, Wang X, et al. The Conserved Non-coding Sequences CNS6 and CNS9 Control Cytokine-Induced Rorc Transcription during T Helper 17 Cell Differentiation. Immunity. 2020;53(3):614–26 e4.

90. Kennedy A, Waters E, Rowshanravan B, Hinze C, Williams C, Janman D, et al. Differences in CD80 and CD86 transendocytosis reveal CD86 as a key target for CTLA-4 immune regulation. Nat Immunol. 2022;23(9):1365–78.

91. Qureshi OS, Zheng Y, Nakamura K, Attridge K, Manzotti C, Schmidt EM, et al. Trans-endocytosis of CD80 and CD86: a molecular basis for the cell-extrinsic function of CTLA-4. Science. 2011;332(6029):600–3.

92. Tekguc M, Wing JB, Osaki M, Long J, and Sakaguchi S. Treg-expressed CTLA-4 depletes CD80/CD86 by trogocytosis, releasing free PD-L1 on antigen-presenting cells. Proc Natl Acad Sci U S A. 2021;118(30).

93. Oderup C, Cederbom L, Makowska A, Cilio CM, and Ivars F. Cytotoxic T lymphocyte antigen-4-dependent down-modulation of costimulatory molecules on dendritic cells in CD4+ CD25+ regulatory T-cell-mediated suppression. Immunology. 2006;118(2):240–9.

