## Supplemental Figures for "Regulatory T cells epigenetically reprogrammed from autoreactive effector T cells mitigate established autoimmunity"

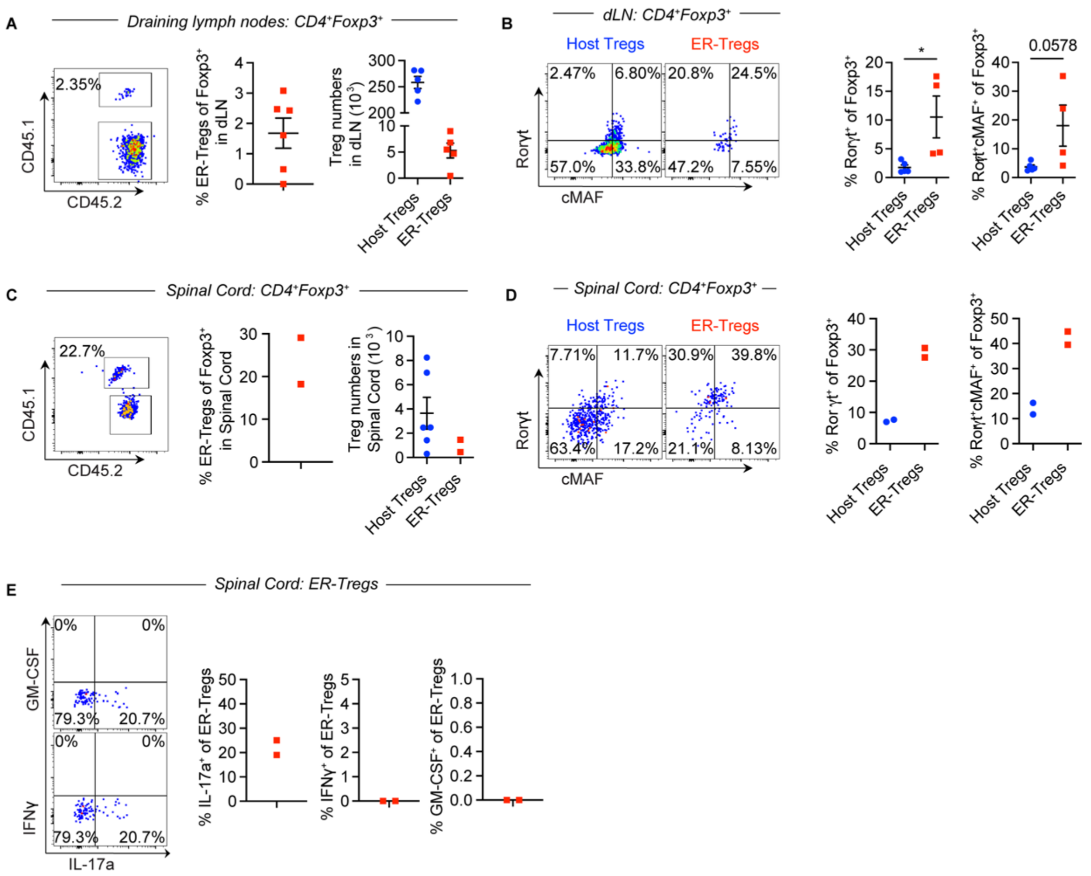


**Supplemental Figure 1. Cellularity and phenotypes of transferred ER-$T_{regs}$ in mice with EAE**

(**A-B**) EAE was induced via MOG/CFA immunization in CD45.2$^+$ mice with or without adoptive transfer of ER-$T_{regs}$ reprogrammed from MOG/CFA-primed CD45.1$^+$*Foxp3$^{Thy1.1}$* CD4$^+$ $T_{eff}$ cells, administered one day prior to immunization. Flow cytometry analyses were conducted at 21 days post-immunization (dpi). n = 6 per group. Data are representative of two independent experiments. (**A**) Flow cytometry analysis of the frequency and total number of host $T_{regs}$ and transferred ER-$T_{regs}$ in the draining lymph nodes. (**B**) Flow cytometry analysis of Rorγt and c-MAF expression in both host $T_{regs}$ and ER-$T_{regs}$ in the draining lymph nodes. (**C-E**) EAE was induced via MOG/CFA immunization in CD45.2$^+$ mice with or without adoptive transfer of ER-$T_{regs}$ reprogrammed from MOG/CFA-primed CD45.1$^+$*Foxp3$^{Thy1.1}$* $T_{eff}$ cells, administered at 11 dpi. Flow cytometry analyses were conducted at 29 dpi. n = 6 per group. Data are representative of two independent experiments. (**C**) Flow cytometry analysis of the frequency of ER-$T_{regs}$ and the total number of host $T_{regs}$ and ER-$T_{regs}$ in the spinal cords. (**D**) Flow cytometry analysis of Rorγt and c-MAF expression in both host $T_{regs}$ and ER-$T_{regs}$ in the spinal cords. (**E**) Flow cytometry analysis of cytokine expression in spinal cord ER-$T_{regs}$. Mean ± SEM. *$p < 0.05$. Unpaired two-sided t-test in (**B**).

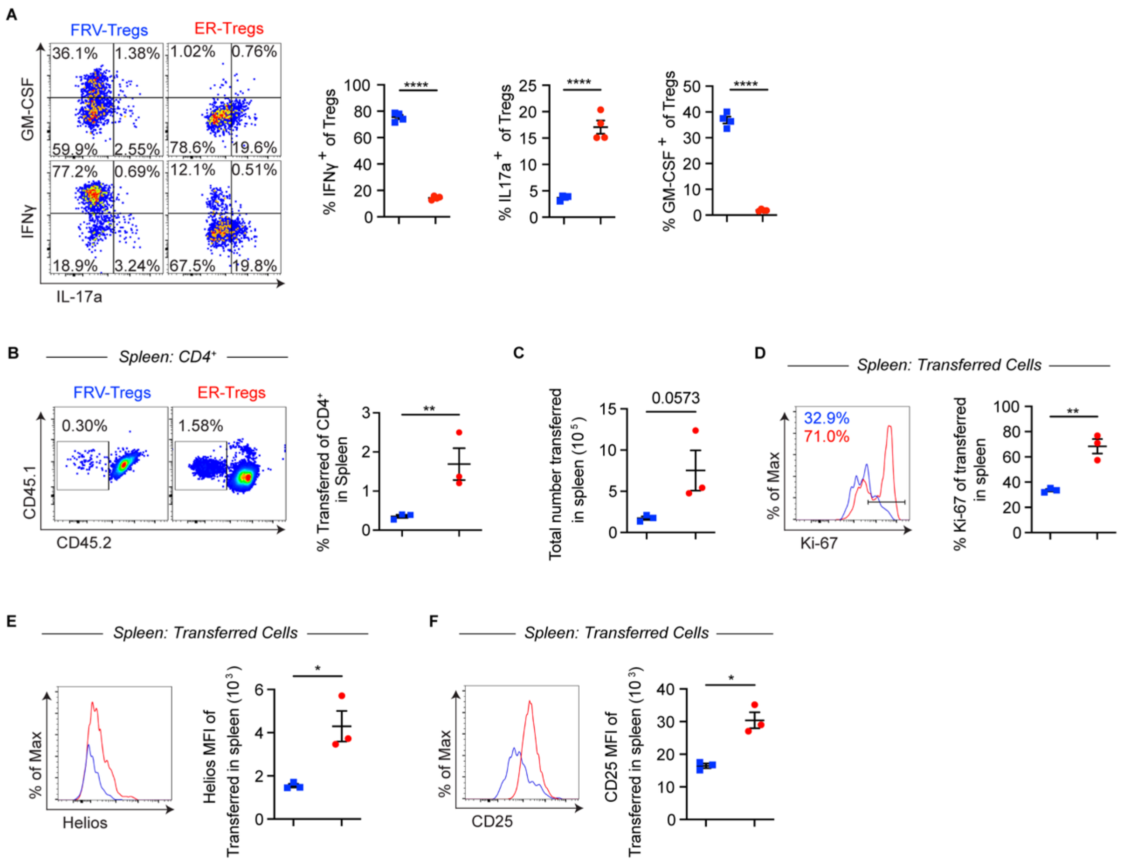


**Supplemental Figure 2. Cellularity, proliferation, and phenotypes of FRV-$T_{regs}$ and ER-$T_{regs}$**

(**A**) $T_{eff}$ cells retrovirally transduced with a Foxp3 overexpression vector (FRV-$T_{regs}$) or ER-$T_{regs}$ were co-cultured with APCs and responder $T_{eff}$ in the presence of MOG for 3 days before flow cytometry analysis of IFNγ, IL-17a, and GM-CSF expression. (**B**-**F**) FRV-$T_{regs}$ or ER-$T_{regs}$ were transferred into CD45 congenically distinct mice. Recipients were immunized with MOG/CFA and cells were analyzed with flow cytometry 4 days after immunization. (**B)** Flow cytometry plots (left) and quantification (right) of the frequencies of transferred cells in the spleens of recipient mice. (**C)** Numbers of transferred cells in the spleens of recipient mice. (**D**) Flow cytometry histograms (left) and quantification (right) of Ki-67 MFI of transferred cells in the spleens of recipient mice. (**E**) Flow cytometry histograms (left) and quantification (right) of Helios MFI of transferred cells in the spleens of recipient mice. (**F**) Flow cytometry histograms (left) and quantification (right) of CD25 MFI of transferred cells in the spleens of recipient mice. Mean ± SEM in. $^*p < 0.05$, $^{**}p < 0.01$, $^{****}p<0.0001$. Unpaired two-sided t-test in (**A**-**F**).

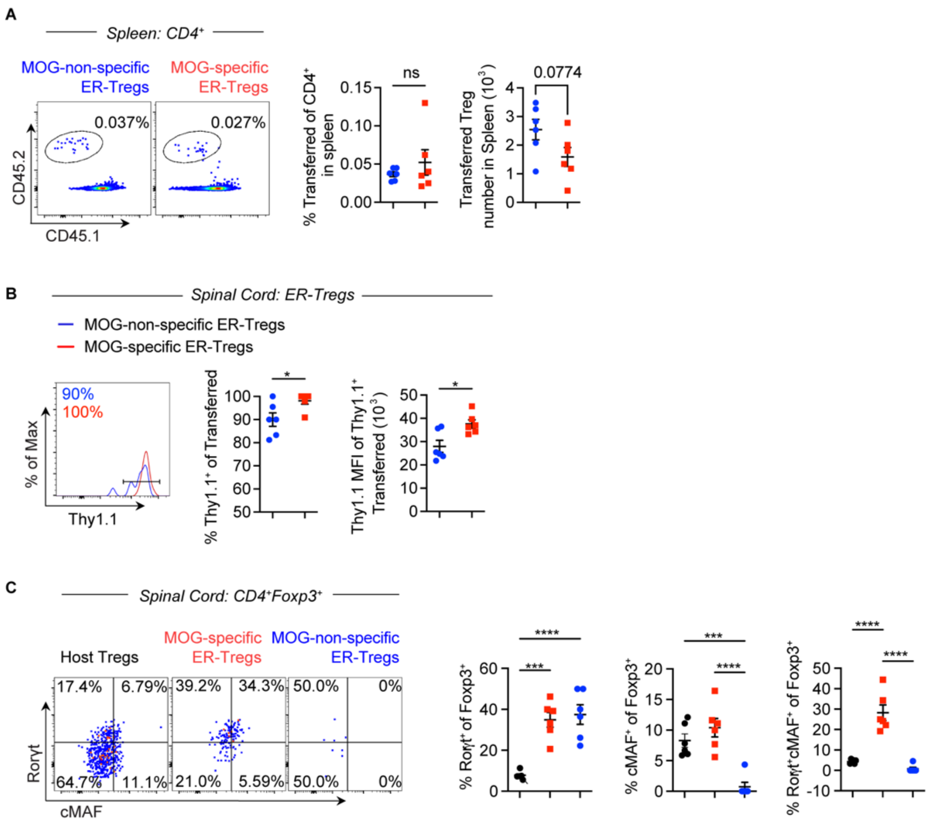


**Supplemental Figure 3. Stability, phenotype, and cellularity of MOG-specific and MOG-nonspecific ER-$T_{regs}$ in EAE**

(**A-C**) EAE was induced via MOG/CFA immunization in $CD45.1^+$ mice with or without adoptive transfer of $CD45.2^+$*$Foxp3^{Thy1.1}$* MOG-specific or MOG-non-specific ER-$T_{regs}$, administered at 11 dpi. Flow cytometry analyses were conducted at 33 dpi. n = 6 per group. (**A**) Flow cytometry analysis of the frequency and total cell number of MOG specific and MOG non-specific ER-$T_{regs}$ in the spleen. (**B**) Flow cytometry analysis of Thy1.1 expression and Thy1.1 MFI in MOG specific vs MOG non-specific ER-$T_{regs}$ in the spinal cord. (**C**) Flow cytometry analysis of Rorγt and c-MAF expression in host $T_{regs}$ and ER-$T_{regs}$ in the spinal cord. Mean ± SEM. *$p<0.05$, ***$p < 0.001$, ****$p < 0.0001$. Unpaired two-sided t-test in (**A**) and (**B**), one-way ANOVA and Holm-Šídák test in (**C**).

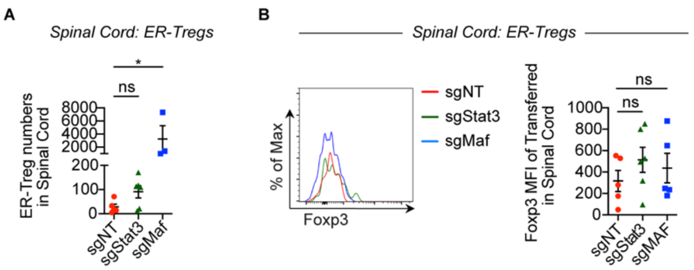


**Supplemental Figure 4. Cellularity and stability of sgNT, sgStat3, and sgMaf transduced ER-T$_{regs}$**

EAE was induced via MOG/CFA immunization in *Rag1*$^{-/-}$ mice one day after adoptive transfer of MOG/CFA-primed CD45.1$^{+}$ CD4$^{+}$ T$_{conv}$ cells with or without co-transfer of CD45.2$^{+}$*Foxp3*$^{Thy1.1}$*R26*$^{Cas9}$ ER-T$_{regs}$ transduced with sgRNA-RV targeting *Stat3* (sgStat3), *Maf* (sgMaf), or a non-targeting sgRNA-RV (sgNT). Flow cytometry analyses were conducted at 22 dpi. n = 6-7 per group. (**A**) Total number of ER-T$_{regs}$ in the spinal cords. (**B**) Flow cytometry histograms (left) and quantification (right) of ER-T$_{reg}$ Foxp3 expression in the spinal cord. Mean ± SEM in (**A-B**). *$p < 0.05$. One-way ANOVA and Holm-Šídák test in (**A-B**).
